# Pyronaridine Protects Against SARS-CoV-2 in Mouse

**DOI:** 10.1101/2021.09.30.462449

**Authors:** Ana C. Puhl, Giovanni F. Gomes, Samara Damasceno, Andre S. Godoy, Gabriela D. Noske, Aline M. Nakamura, Victor O. Gawriljuk, Rafaela S. Fernandes, Natalia Monakhova, Olga Riabova, Thomas R. Lane, Vadim Makarov, Flavio P. Veras, Sabrina S. Batah, Alexandre T. Fabro, Glaucius Oliva, Fernando Q. Cunha, José C. Alves-Filho, Thiago M. Cunha, Sean Ekins

## Abstract

There are currently relatively few small-molecule antiviral drugs that are either approved or emergency approved for use against SARS-CoV-2. One of these is remdesivir, which was originally repurposed from its use against Ebola and functions by causing early RNA chain termination. We used this as justification to evaluate three molecules we had previously identified computationally with antiviral activity against Ebola and Marburg. Out of these we previously identified pyronaridine, which inhibited the SARS-CoV-2 replication in A549-ACE2 cells. Herein, the *in vivo* efficacy of pyronaridine has now been assessed in a K18-hACE transgenic mouse model of COVID-19. Pyronaridine treatment demonstrated a statistically significant reduction of viral load in the lungs of SARS CoV-2 infected mice. Furthermore, the pyronaridine treated group reduced lung pathology, which was also associated with significant reduction in the levels of pro-inflammatory cytokines/chemokine and cell infiltration. Notably, pyronaridine inhibited the viral PL^pro^ activity *in vitro* (IC_50_ of 1.8 µM) without any effect on M^pro^, indicating a possible molecular mechanism involved in its ability to inhibit SARS-CoV-2 replication. Interestingly, pyronaridine also selectively inhibits the host kinase CAMK1 (IC_50_ of 2.4 µM). We have also generated several pyronaridine analogs to assist in understanding the structure activity relationship for PL^pro^ inhibition. Our results indicate that pyronaridine is a potential therapeutic candidate for COVID-19.

**One sentence summary:** There is currently intense interest in discovering small molecules with direct antiviral activity against the severe acute respiratory syndrome coronavirus 2 (SARS-Cov-2). Pyronaridine, an antiviral drug with in vitro activity against Ebola, Marburg and SARS-CoV-2 has now statistically significantly reduced the viral load in mice along with IL-6, TNF-α, and IFN-β ultimately demonstrating a protective effect against lung damage by infection to provide a new potential treatment for testing clinically.

## Introduction

At the time of writing, we are in the midst of a major a global health crisis caused by the virus Severe Acute Respiratory Syndrome Coronavirus 2 (SARS-CoV-2) that was originally reported in Wuhan, China in late 2019 (*1, 2*). Infection with this virus leads to extensive morbidity, mortality and a very broad range of clinical symptoms such as cough, loss of smell and taste, respiratory distress, pneumonia and extrapulmonary events characterized by a sepsis-like disease collectively called 2019 coronavirus disease (COVID-19) (*3*). In the USA, there are currently three vaccines available, one of which has recently obtained full approval from the FDA to protect against SARS-CoV-2 (*4–6*). There are however few small-molecule drugs approved for COVID-19 (*7*) including remdesivir (*8*), which originally demonstrated activity in Vero cells (*9, 10*), human epithelial cells and in Calu-3 cells (*10*) infected with SARS-CoV-2 prior to clinical testing. Remdesivir represents a repurposed drug which was originally developed for Hepatitis C virus but was then repurposed for treating Ebola and has since reached clinical trials (*11*). We therefore hypothesized that other drugs that were effective against Ebola might also be prioritized for evaluation *in vitro* against SARS-CoV-2. Previously, we had used a machine-learning model to identify tilorone, quinacrine and pyronaridine tetraphosphate(*12*) for testing against Ebola virus (EBOV) and subsequently these three inhibited EBOV and Marburg *in vitro* as well as demonstrating significant efficacy in the mouse-adapted EBOV (ma-EBOV) model (*13–15*). All of these molecules were identified as lysosomotropic, a characteristic that suggests these could be possible entry inhibitors (*16*). Pyronaridine tetraphosphate is used as an antimalarial in several countries as part of a combination therapy with artesunate (Pyramax). Pyronaridine alone also demonstrated significant activity in the guinea pig-adapted model of EBOV infection (*17*). We and others (*18–20*) have recently shown that these compounds possess *in vitro* activity against SARS-CoV-2 and tilorone and pyronaridine are in clinical trials, the latter in combination with artesunate. The C_max_ data for pyronaridine in our previous mouse pharmacokinetics studies (i.p. dosing) suggests that plasma levels that are above the average IC_50_ observed for SARS-CoV-2 inhibition *in vitro* (*13*) can be reached with dosing well below the maximum tolerated dose. Pyronaridine also has excellent *in vitro* ADME properties with a long half-life that makes a single dose treatment possible (*13, 18*). We now expand on our earlier *in vitro* characterization of pyronaridine (*18*) by assessing the *in vivo* efficacy in a mouse model of COVID-19. Finally, in an attempt to further explore molecular mechanisms, we tested the activity of pyronaridine *in vitro* against viral and host targets.

### *In vivo* efficacy of Pyronaridine in a mouse model of COVID-19

*In vivo* efficacy was assessed in K-18-hACE2 mouse model of COVID-19 (*21–23*). Pyronaridine (75 mg/kg, i.p) (*13*) was administered 1 h prior to infection. Mice that were given pyronaridine, received a single treatment. On the third day post-infection, mice were euthanized and lung viral load, cytokine levels and histopathology were evaluated (Figure 1A). In all groups tested, mice lost weight compared to uninfected animals that received only vehicle formulation (Figure 1B). Lung viral load was evaluated by RT-qPCR and the pyronaridine treated group showed a statistically significant decrease in the lung viral load (Figure 1C). Moreover, reduced levels of INF-1β were observed in infected mice, and pyronaridine restored the levels of INF-1β close to that found in uninfected animals (Figure 1D). Interestingly, the increased levels of IL-6 found in the infected untreated group were not observed in animals treated with pyronaridine (Figure 1E). In addition, pyronaridine reduced the high levels of CXCL1 and CCL4 observed in infected animals (Figure 1D). Pyronaridine did not however reduce the high levels of IL-10, TNF-α, CCL2, and CCL3 (Figure 1F, G, K, L, M) below the elevated levels found in the infected mice.

**Figure 1:**
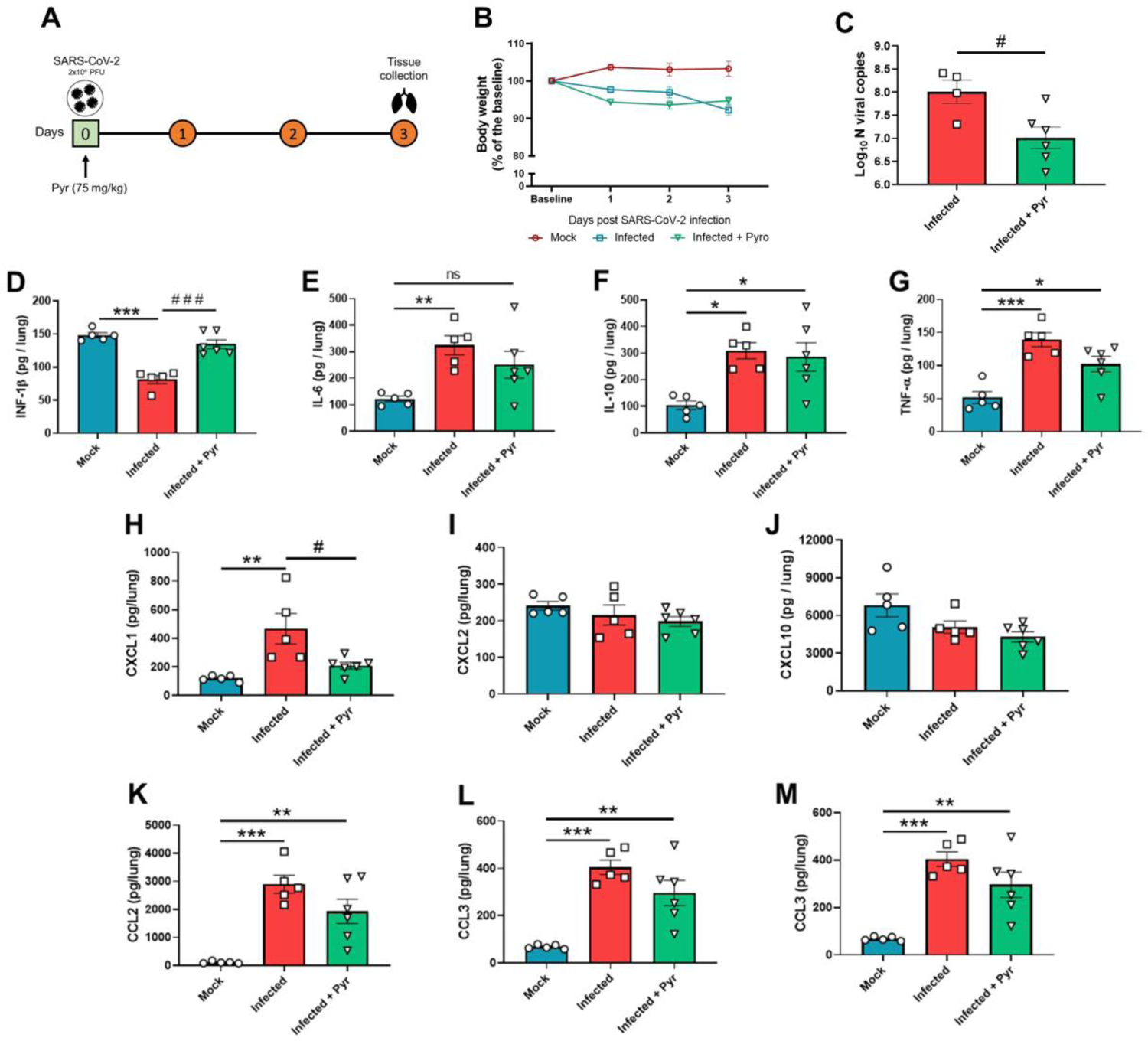
*In vivo* efficacy of pyronaridine in a mouse model of COVID-19. A) Experimental timeline: K18-hACE2 mice were infected with SARS-CoV-2 (2 × 10^4^ PFU/40 µL saline, intranasal) or mock. One group of mice was treated with pyronaridine (75 mg/kg i.p.) 1 h before virus inoculation. B) Body weight was evaluated daily. C) At 3 DPI, mice were euthanized and the lung viral load, and D-M) Lung cytokines and chemokines levels were determined. * p<0.05, ** p<0.01, and *** p<0.001 as compared with mock group after one-way ANOVA followed by Tukey post-hoc test. # p<0.05 as compared with infected group after one-way ANOVA followed by Tukey post-hoc test. Pyr – Pyronaridine.

To determine the severity of lung damage, histological examination of hematoxylin and eosin (H&E) stained lung tissue was performed. Infected, untreated mice showed severe pathological changes with inflammatory cell infiltrates. In contrast, pyronaridine-treated animals exhibited improved morphology and milder infiltration (Figure 2A). Histological observations were confirmed by quantitative morphometric analysis of the H&E-stained slides showing a statistically significant reduction in inflammation (Figure 2B).

**Figure 2:**
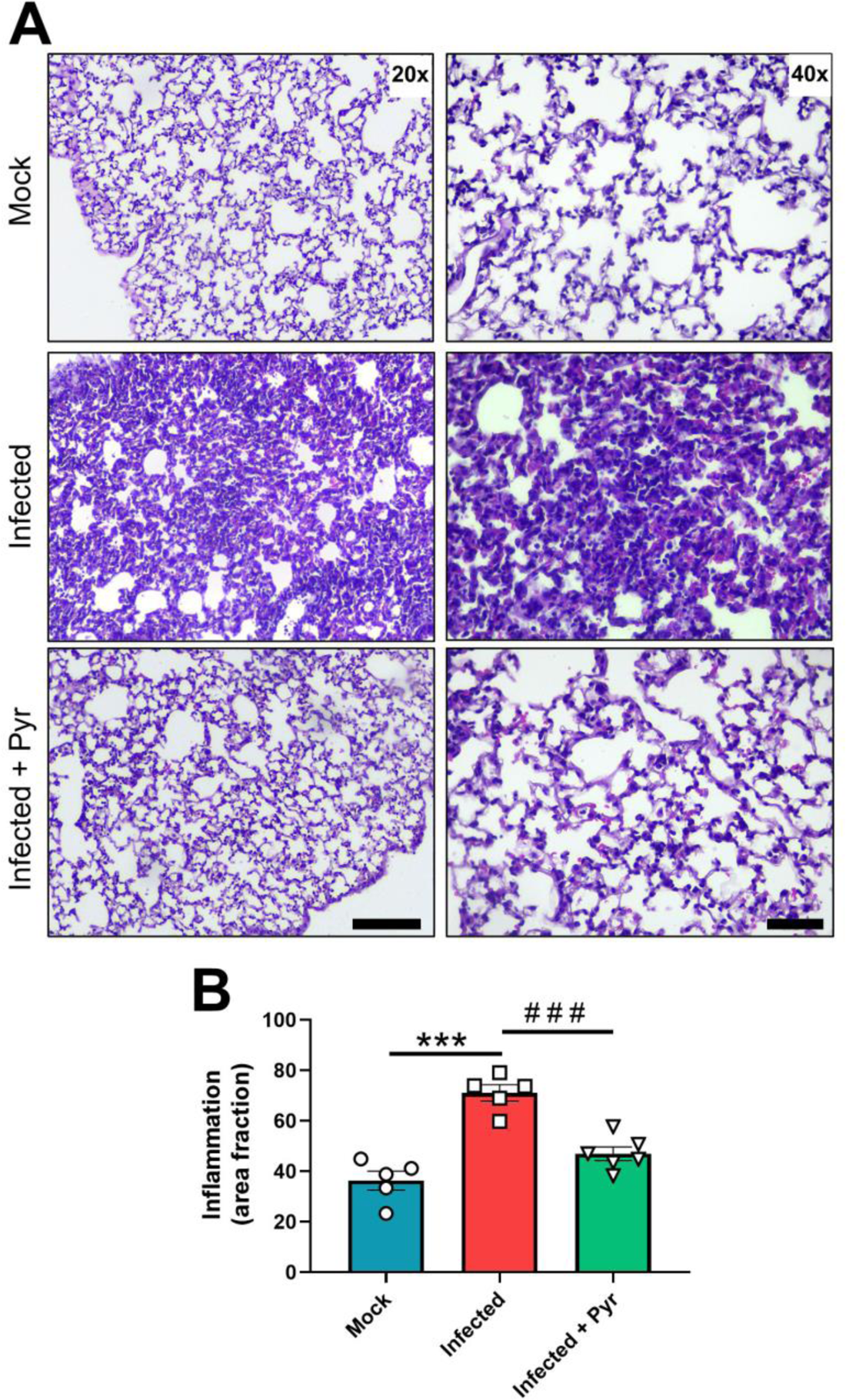
Lung histopathological analyses of COVID-19 mice treated with pyronaridine. K18-hACE2 mice were infected with SARS-CoV-2 (2 × 10^4^ PFU/40 µL, intranasal) or mock. One group of mice was treated with pyronaridine (75 mg/kg i.p.) 1 h before virus inoculation. At 3 DPI, mice were euthanized, and the lungs were harvested and processed for histopathological analyses. A - Representative images of lung slices stained with hematoxylin and eosin (H&E). B-Quantitative morphometric analyses. *** p<0.001 as compared with mock group after one-way ANOVA followed by Tukey post-hoc test. # p<0.05 and ## p<0.01 as compared with infected group after one-way ANOVA followed by Tukey post-hoc test. Pyr – Pyronaridine. Scale bars: 20X = 125 µm; 40X = 50 µm.

### Pyronaridine potentially targets SARS-CoV-2 PL^pro^ but not M^pro^

SARS-CoV-2 proteases M^pro^ and PL^pro^ are essential for viral replication and have been widely studied for the discovery of new direct acting antivirals (*24–26*). Pyronaridine was therefore tested against both SARS-CoV-2 recombinant PL^pro^ and M^pro^ through FRET-based *in vitro* assays. Pyronaridine inhibited PL^pro^ with an IC_50_ of 1.86 ±0.58 µM (Figure 3A) but did not show any appreciable activity against M^pro^ at 20 µM (data not shown). Additional analogs of pyronaridine were also synthesized and tested against PL^pro^. The analogs 12126038, 12126039 and 12126040 showed similar inhibitory activity when compared with pyronaridine (as well as that reported for GRL0617 (*27*)), indicating that the aminophenol moiety together with pyrrole or tertiary amine substitutions at the meta position are tolerated for PL^pro^ inhibition (Table 1, Figure 3A). The deletion of these groups in analogs 10326099, 12126035, 12126036, 12126037 and 12126072 caused complete abolishment of the inhibitory activity of the series (Table 1). The PL^pro^ active site contains four subsites for peptide recognition, with a strong preference for positively charged amino acids at P3 and P4 subsites (*24–26*). In our molecular docking studies (Figure 3B), the positions where the 2,6-bis[(pyrrolidin-1-yl)methyl]phenol moiety of pyronaridine were docked at the P4 subsite showed a far higher score than the second highest cluster group (Glide score of −5.052 and HADDOCK score of −65.6). These positively charged amino-groups would likely satisfy the negatively charged cavity that forms the P4 subsite, forming hydrogen bounds with Asp164 and π-stacking with Tyr26 (Figure 3C). Compounds 12126038 and 12126039 also showed very similar poses after docking, indicating binding to the P4 subsite (Figure 3B). The primary docking poses from Glide for all compounds are provided in Fig. S1. These results indicate that the *in vitro* and *in vivo* efficacy of pyronaridine may be related to PL^pro^ inhibition.

**Figure 3.**
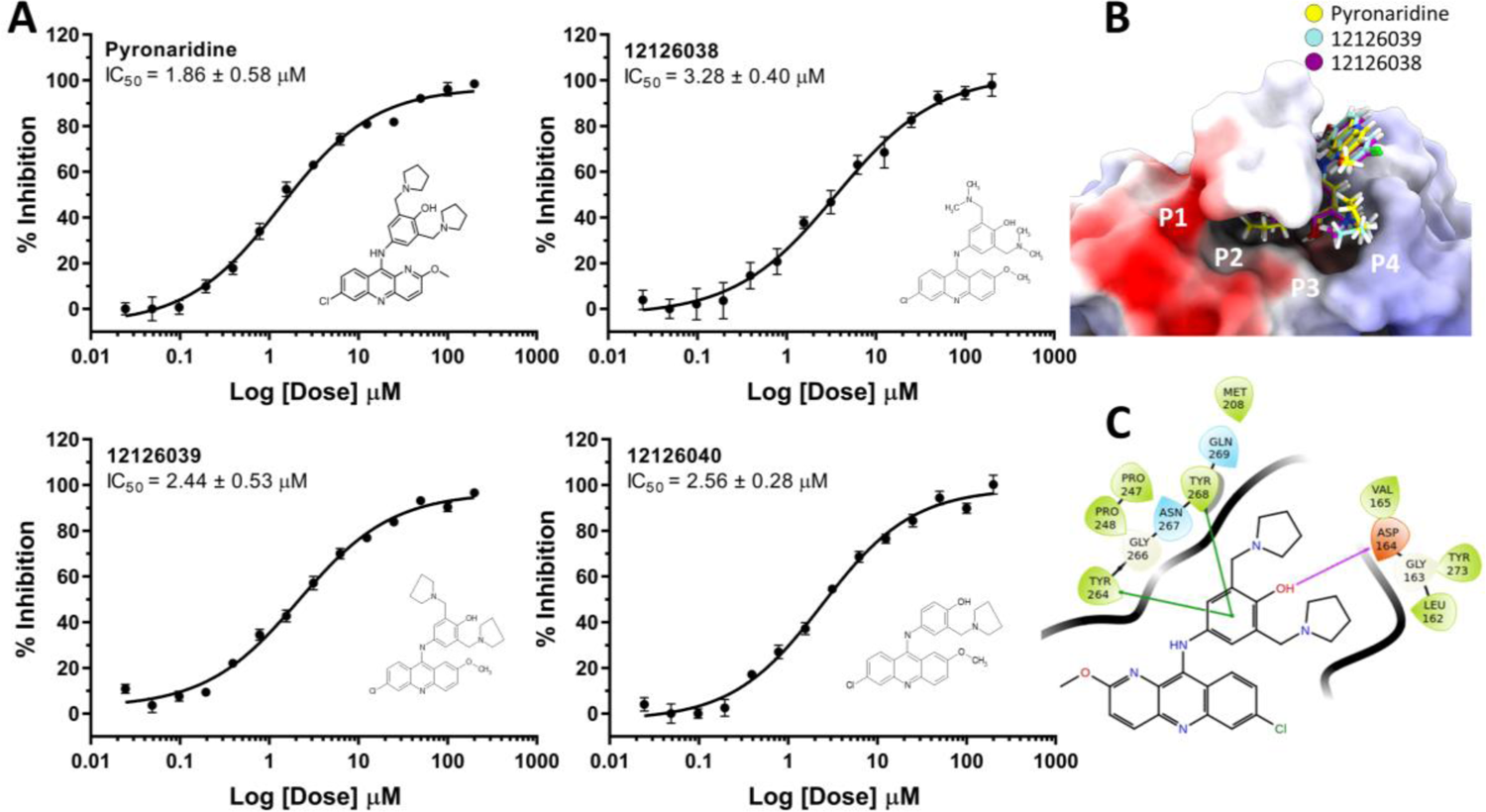
A. Dose-response curves of pyronaridine and active analogs against SARS-CoV-2 PL^pro^ B. Docking pose of pyronaridine (yellow) and analogs 12126039 (blue) and 12126038 (purple) in PL^pro^ active site (PDB 7JRN). Electrostatic potential projected as surface charge. C. 2D ligand interaction diagram. Hydrogen bonds are depicted in pink, while pi-stackings are depicted in green.

**Table 1.**
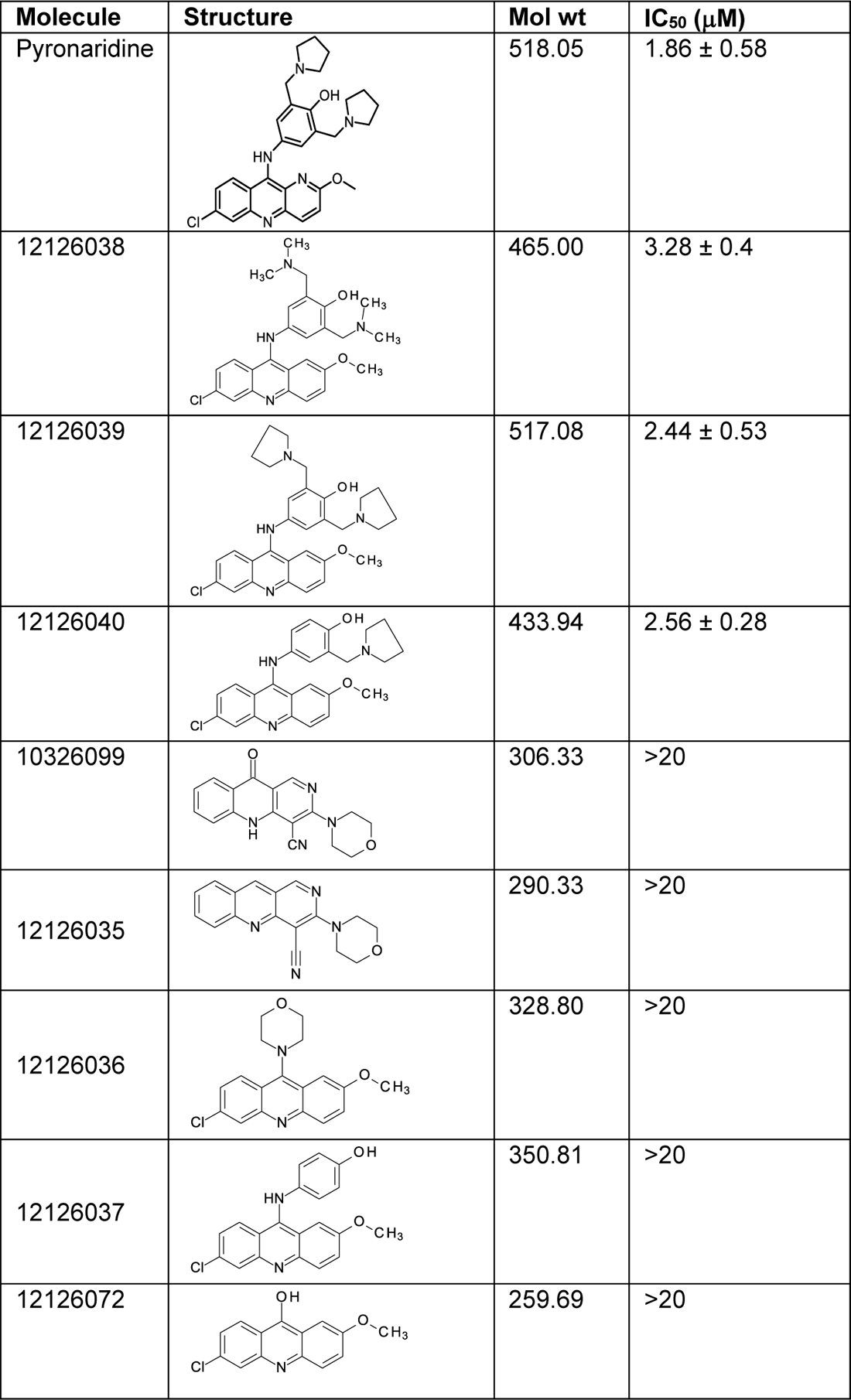
Pl^pro^ IC_50_ inhibition data for pyronaridine analogs.

| Molecule | Structure | Mol wt | IC<sub>50</sub> (μM) |
| --- | --- | --- | --- |
| Pyronaridine |  | 518.05 | 1.86 ± 0.58 |
| 12126038 |  | 465.00 | 3.28 ± 0.4 |
| 12126039 |  | 517.08 | 2.44 ± 0.53 |
| 12126040 |  | 433.94 | 2.56 ± 0.28 |
| 10326099 |  | 306.33 | >20 |
| 12126035 |  | 290.33 | >20 |
| 12126036 |  | 328.80 | >20 |
| 12126037 |  | 350.81 | >20 |
| 12126072 |  | 259.69 | >20 |

### Kinase activity profiling

Due to the possible effect of pyronaridine on cytokines (Figure 1) we have also assessed the effect on host targets, as SARS-CoV-2 can cause an imbalance in the immune system that may result in a cytokine storm(*28*) as well as leading to acute respiratory distress syndrome (ARDS), coagulation disorders, and eventually multiple organ failure (*28, 29*). Hence targeting the cytokine storm to address hyperinflammation represents another approach to the treatment of COVID-19 patients (*30–33*). In this regard, we have explored the effect of pyronaridine on human kinases which are responsible for host cell signaling (*34, 35*). Screening of pyronaridine (tested at 1 µM) against 485 kinases identified only 2 as having mean percent inhibition greater than 30% including CAMK1 (35%) and MELK (31%) (Table S1). Subsequently, the IC_50_ was determined for CAMK1 (2.4 µM) (Figure 4).

**Figure 4:**
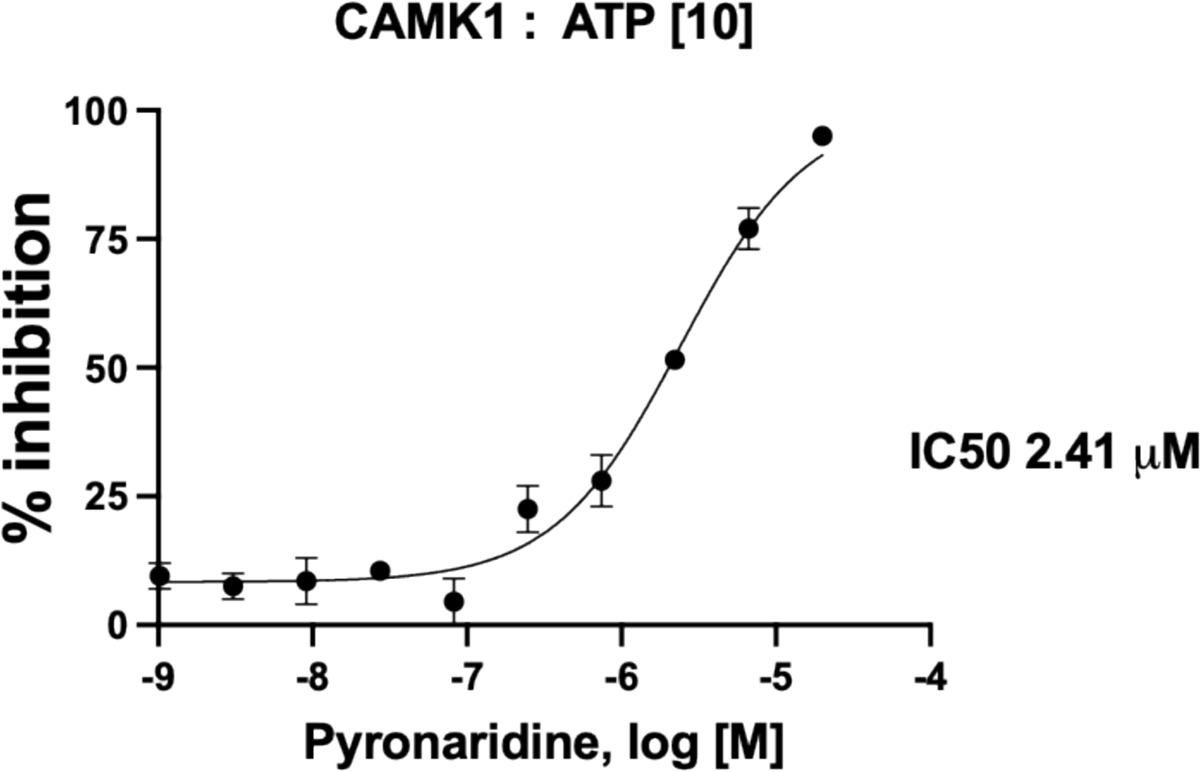
Pyronaridine CAMK1 dose response.

## Discussion

Even with vaccines becoming widely available in many countries COVID-19 continues to exact a very heavy toll on those that are unvaccinated. We are in a race against time before the virus mutates and vaccines lose their effectiveness. There is therefore an urgent need for new antivirals and in particular small-molecule treatments that are orally delivered and can be used outside of a hospital setting. Finding, developing, and progressing small molecules to the clinic is generally a slow and expensive process (*36*), hence drug repurposing has been attempted by many groups to speed this up (either experimentally or computationally (*37*)) by identifying already approved or clinical stage candidates used for other applications or quickly follow up the few molecules that are being used already. The traditional prioritization of compounds *in vitro* before animal models and then humans is still repeated and so far with few successes with many molecules not demonstrating efficacy *in vivo* (*38, 39*).

Our understanding of the antiviral mechanism of pyronaridine previously shown to inhibit the Ebola virus *in vitro* and *in vivo* (*13*) via binding to the viral glycoprotein(*16*) as well as through its potent lysosomotropic activity (*40*) and now the *in vitro* activity against SARS-CoV-2 (*18*) is also further expanded. Pyronaridine was previously identified with *in vitro* activity against SARS-CoV-2 in A549-ACE2 cells that was on a par with remdesivir in this cell line (*18*). In the current study, we have demonstrated that pyronaridine also has antiviral activity against SARS-CoV-2 *in vivo.* A single prophylactic dose of pyronaridine (75 mg/kg i.p) reduced the viral load in the lung of infected mice 3 days post-infection. *In vitro* assays suggest that pyronaridine possesses a direct antiviral effect showing activity against PL^pro^ (IC_50_ 1.86 µM, Figure 3), but did not inhibit SARS-CoV-2 M^pro^. Kinase profiling, resulted in determination of IC_50_ for CAMK1 (IC_50_ 2.4 µM, Figure 4).

A single dose (75 mg/kg i.p.) of pyronaridine has an elimination half-life of 146 h in mice (*13*), comparable to the reported half-life of pyronaridine in humans of 195-251 h (*41, 42*). A study of pyronaridine as a single oral dose (400 mg) given to a healthy volunteer found a C_max_ in plasma of 495.8 ng/ml at a T_max_ of 0.5 h (*41, 43*), which gives a concentration close to 1 µM. In our study, pyronaridine inhibits Pl^pro^ at 1.86 µM and CAMK1 at 2.4 µM, which is very close to the C_max_ of 1 µM. Pyronaridine preferentially associates with blood cells and is highly plasma protein bound (*43*), which suggests that it may not reach the unbound concentration necessary to have an effect on Pl^pro^ and CAMK1 at these doses. Activation of CAMK1 has been reported to negatively impact HBV biosynthesis (*44, 45*), however the mechanism has not been elucidated yet and the possible role of CAMK1 on SARS-CoV-2 infection remains to be investigated.

Several studies have shown a cytokine and chemokine storm as important during SARS-CoV-2 infection in patients with COVID-19, including PDGF, VEGF (*46*), IL-6 (*47, 48*), IL-8, IL-10 (*48*), TNF-α (*48*) and IFN-α and -β (*49–53*). Furthermore, an impaired type I interferon response has already been observed in COVID-19 (*54*), which is followed by increased circulating levels of IL-6 and TNF-α. Many viruses, including SARS-CoV-2, subvert the immune system by inhibiting the production of interferons (INF), an important family of antiviral mediators (*55–58*), which results in an impaired antiviral defense and increased viral replication and infection. Here, we have demonstrated that pyronaridine restored the levels of INF-1β in infected mice. Moreover, as already discussed, this effect was associated with a reduced viral load after the treatment with pyronaridine. Notably, pyronaridine also reverted the altered levels of IL-6 and CXCL1 observed in infected mice. In addition to biochemical changes, histopathological findings are also observed in the lung of patients (*59–61*) and in other experimental models of COVID-19 (*21–23*). In the present study, control animals infected with SARS-CoV-2 presented pronounced inflammatory cell infiltration and the histological examination of mouse lungs for those treated with pyronaridine showed milder infiltration, appearing comparable to that of uninfected mice. Thus, pyronaridine appears to have both antiviral and immunomodulatory effects in this experimental model of COVID-19 as used in the present study.

In summary, our present study provided additional data on the efficacy of pyronaridine against SARS-CoV-2 infection as well as highlighting reduced lung pathology and inflammation in a mouse model of COVID-19. Furthermore, we have shown that pyronaridine may target Pl^pro^ as well as CAMK1. There are few inhibitors of CAMK1 that have been identified to date (such as Barettin (*62*) or pyridine amides (*63*)) which has a role in inflammation targeting IL-10 (*62*). Previous *in vitro* work has shown that inhibiting CAMK1 in cells reduces IL-10, the master anti-inflammatory interleukin (*64*). In the present study there is not a significant difference in the IL-10 levels between the untreated and pyronaridine-treated infected groups so it seems unlikely that CAMK1 inhibition would be involved in the mechanism of action of inhibition of SAR-CoV-2. In conclusion, we propose that pyronaridine could be used alone as a potential therapeutic candidate for COVID-19. Finally, with the emerging virulence of novel SARS-CoV-2 strains, identifying repurposed drugs with novel mechanisms of action and whose antiviral activity translates from *in vitro* to *in vivo* are rare (*39*), and may lead to new treatments as well as their further optimization.

## ACKNOWLEDGMENTS

The authors would like to kindly acknowledge their many collaborators around the world who have assisted in our various COVID-19 projects. The authors gratefully acknowledge the technical assistance of Marcella Daruge Grando, Ieda Regina dos Santos, Juliana Trench Abumansur, and Felipe Souza.

## Author contributions

A.C.P, T.M.C, S.E. conceived and codirected the study. A.C.P., G.F.G, A.S.G., V.M., G.O., F.Q.C., J.C.A.F, T.C designed the experiments; A.C.P, S.D., G.D.N., A.M.N, V.O.G, R.S.F, N.M., O.R, S.S.B and A.T.F performed *in vitro* experiments. G.F.G, S.D. F.P.V, performed *in vivo* experiments. A.C.P, S.E., G.F.G, A.S.G. and T.M.C, and T.L. wrote the manuscript. All authors read and accept the manuscript.

## Materials and Methods

### Chemicals and reagents

Pyronaridine tetraphosphate [4-[(7-Chloro-2-methoxybenzo[b][1,5]naphthyridin-10-yl)amino]-2,6-bis(1-pyrrolidinylmethyl)phenol phosphate (1:4)] (*12*) was purchased from BOC Sciences (Shirley NY). The purity of this compound was greater than 95%. For pyronaridine analogs, ^1^H and ^13^C Spectra were measured on Bruker AC-300 (300 MHz, ^1^H) or Bruker AC-200 (50 MHz, ^13^C). Chemical shifts were measured in DMSO-d_6_or CDCl_3_, using tetramethylsilane as an internal standard, and reported as units (ppm) values. The following abbreviations are used to indicate the multiplicity: s, singlet; d, doublet; t, triplet; m, multiplet; dd, doublet of doublets; brs, broad singlet; brm, broad multiplet. The purity of the final compounds was analyzed on Agilent 1290 Infinity II HPLC system coupled to Agilent 6460 triple-quadrupole mass spectrometer equipped with an electrospray ionization source. The chromatographic separation was carried out on Agilent Eclipse Plus C18 RRHD column (2.1 × 50 mm, 1.8 µm) at 40 °C, sample injection volume was 0.2 µL.The mobile phase comprising 0.1 % formic acid / water (A), and 0.1 %formic acid and 85 % acetonitrile / water (B) was programmed with gradient elution (0.0-3.0 min, 60 % B; 3.0-4.0 min, 60 % to 97 % B; 4.0-6.0 min, 97 % B; 6.0-6.1 min, 97 % to 60 % B) at a flow rate of 0.4 mL/min. The mass spectrometric detection was operated in positive ion mode. Optimal parameters were: capillary voltages of 3500 V, a nebulizer pressure of 35 psi, a gas temperature of 350 °C, a gas flow rate of 12 L/min. All final compounds are > 95 % pure. Melting points were determined on Electrothermal 9001 (10°C per min) and are uncorrected. Merck KGaAsilica gel 60 F_254_ plates were used for analytical thin-layer chromatography. Column chromatography was performed on Merck silica gel 60 (70-230 mesh). Yields refer to purified products and are not optimized.

The molecules were synthesized according to Scheme 1 or Scheme 2 and the specific methods and analytical results are described in the Supplemental Methods.

### Test Article Preparation

Dose formulation for pyronaridine was prepared as previously described (*13*) under yellow light by mixing the appropriate amount of pyronaridine in melted Kolliphor HS 15 (Solutol) (20% final volume) using a vortex mixer for 30 s. The remaining sterile water (Gibco) was added, and the formulations were mixed using a vortex mixer for 30 sec – 5 min until the compound was visually dissolved and then sonication for 25 min. The final 20% Kolliphor HS 15 dose formulations was observed to be a clear, reddish solutions.

### Mouse studies Ethical approval

All the experimental procedures were performed in accordance with the guide for the use of laboratory animals of the University of Sao Paulo and approved by the institutional ethics committee under the protocol number 105/2021.

### SARS-CoV-2 isolate

SARS-CoV-2 was isolated from a COVID-19 positive-tested patient. The virus was propagated and titrated in Vero E6 cells in a biosafety level 3 laboratory (BSL3) at the Ribeirão Preto Medical school (Ribeirão Preto, Brazil). Cells were cultured in DMEM medium supplemented with 10% fetal bovine serum (FBS) and antibiotic/antimycotic (Penicillin 10,000 U/mL; Streptomycin 10,000 μg/mL). The viral inoculum was added to Vero cells in DMEM 2% FBS and incubated at 37 °C with 5% CO_2_ for 48 h. The cytopathogenic effect (CPE) was observed under a microscope. Cell monolayer was collected, and the supernatant was stored in −70 °C. Virus titration was made by the plaque-forming units (PFU).

### K18-hACE2 mice

To evaluate the effects of Pyronaridine *in vivo*, we infected the K18-hACE2 humanized mice (B6.Cg-Tg(K18-ACE2)2Prlmn/J)(*21, 65, 66*). K18-hACE2 mice were obtained from The Jackson Laboratory and were breed in the Centro de Criação de Animais Especiais (Ribeirão Preto Medical School/University of São Paulo). This mouse model for SARS-CoV-2-induced disease has been used as it presents clinical signs, and biochemical and histopathological changes compatible with the human disease(*65–71*). Mice had access to water and food *ad libitum*. For the experimental infection, animals were transferred to the BSL3 facility.

### SARS-CoV-2 experimental infection and treatments

Female K18-hACE2 mice, 8-week-old, were infected with 2×10^4^ PFU of SARS-CoV-2 (in 40 µL) by intranasal route. Uninfected mice (N = 5) were inoculated with equal volume of PBS (Phosphate buffered saline - 137 mM NaCl, 2.7 mM KCl, 10 mM Na_2_HPO_4_, 1.8 mM KH_2_PO_4_; pH 7.4). On the day of infection, 1 h before virus inoculation, animals were treated with Pyronaridine (75 mg/kg, i.p.) (n = 6). Five infected animals remained untreated. Body weight was evaluated on the baseline and on all the days post-infection. On the day 3 post-infection animals were humanely euthanized, and lungs were collected. Right lung was collected, harvested, and homogenized in PBS with steel glass beads. The homogenate was added to TRIzol® (Invitrogen, CA, EUA) reagent (1:1), for posterior viral titration via RT-qPCR, or to lysis buffer (1:1), for ELISA assay, and stored at −70 °C. The left lung was collected in paraformaldehyde (PFA 4%) for posterior histological assessment.

### Absolute viral copies quantification

Total RNA from the right lungs were obtained using the Trizol® (Invitrogen, CA, EUA) method and quantified using NanoDrop One/Onec (ThermoFisher Scientific, USA). A total of 800 ng of RNA was used to synthesize cDNAusing the High-Capacity cDNA Reverse Transcription kit (Applied Biosystems, Foster City, CA, USA), following the manufacturer’s protocol. The determination of the absolute number of viral copies was made by a Taqman real-time qPCR assay with the ad of the StepOne^TM^ Real-Time PCR System (Applied Biosystems, Foster City, CA, USA). A standard curve was generated in order to obtain the exact number of copies in the tested sample, using an amplicon containing 944 bp cloned from a plasmid (PTZ57R/T CloneJet^TM^ Cloning Kit Thermo Fisher®), starting in the nucleotide 14 of the gene N. To quantify the number of copies, a serial dilution of the plasmid in the proportion of 1:10 was performed. Commercial primers and probes for the N1 gene and RNAse P (endogenous control) were used for the quantification (2019-nCov CDC EUA Kit, IDT), following the CDC’s instructions.

### ELISA assay

Lung homogenate was added to RIPA buffer (Radioimmunoprecipitation assay buffer – 150 mM NaCl, 1% NP-40 or Triton X-100, 0.5% sodium deoxycholate, 0.1% SDS, 50 mM Tris; pH 8.0) in proportion of 1:1, and then centrifuged at 10,000 x *g* at 4 °C for 10 min. Supernatant was collected and stored in −70 °C until use. The Sandwich ELISA method was performed to detect the concentration of cytokines and chemokines using kits from R&D Systems (DuoSet), according to the manufacturer’s protocols. The following targets were evaluated: IL-6, IL-10, TNF-α, INF-1β, CCL2, CCL3, CCL4, CXCL1, CXCL2, and CXCL10.

### Lung histopathological process and analyses

Five micrometer lung slices were submitted to Hematoxylin and Eosin staining. A total of 10 photomicrographs in 40X magnification per animal were randomly obtained using a microscope Novel (Novel L3000 LED, China) coupled to a HDI camera for images capture. The total septal area and total area were analyzed with the aid of the Pro Plus 7 software (Media Cybernetics, Inc., MD, USA). Morphometric analysis was performed in accordance with the protocol established by the American Thoracic Society and European Thoracic Society (ATS/ERS) (*72*).

### Synthesis of pyronaridine analogs

All reagents and solvents were purchased from commercial suppliers and used without further purification. ^1^H and ^13^C Spectra were measured on Bruker AC-300 (300 MHz, ^1^H) or Bruker AC-200 (50 MHz, ^13^C). Chemical shifts were measured in DMSO-d_6_or CDCl_3_, using tetramethylsilane as an internal standard, and reported as units (ppm) values. The following abbreviations are used to indicate the multiplicity: s, singlet; d, doublet; t, triplet; m, multiplet; dd, doublet of doublets; brs, broad singlet; brm, broad multiplet.

The purity of the final compounds was analyzed on Agilent 1290 Infinity II HPLC system coupled to Agilent 6460 triple-quadrupole mass spectrometer equipped with an electrospray ionization source. The chromatographic separation was carried out on Agilent Eclipse Plus C18 RRHD column (2.1 × 50 mm, 1.8 µm) at 40 °C, sample injection volume was 0.2 µL.The mobile phase comprising 0.1 % formic acid / water (A), and 0.1 %formic acid and 85 % acetonitrile / water (B) was programmed with gradient elution (0.0-3.0 min, 60 % B; 3.0-4.0 min, 60 % to 97 % B; 4.0-6.0 min, 97 % B; 6.0-6.1 min, 97 % to 60 % B) at a flow rate of 0.4 mL/min. The mass spectrometric detection was operated in positive ion mode. Optimal parameters were: capillary voltages of 3500 V, a nebulizer pressure of 35 psi, a gas temperature of 350 °C, a gas flow rate of 12 L/min. All final compounds are > 95 % pure. Melting points were determined on Electrothermal 9001 (10°C per min) and are uncorrected. Merck KGaAsilica gel 60 F_254_ plates were used for analytical thin-layer chromatography. Column chromatography was performed on Merck silica gel 60 (70-230 mesh). Yields refer to purified products and are not optimized.

The molecules were synthesized according to Scheme 1 or Scheme 2 and the specific methods and analytical results are described in the Supplemental Methods.

### Molecular docking

Molecular docking was performed using Glide software (Schrödinger, 2017, USA) and the SARS-CoV-2 PL^pro^ X-ray structure was downloaded from the Protein Data Bank (PDBid 7JRN). Prior to docking, ligand and protein were prepared with LigPrep (Schrödinger, 2017, USA) and Protein Preparation Wizard (Schrödinger, 2017, USA) using default parameters. Different grids were generated with Receptor Grid Generation at possible sites founded in SiteMap (Schrödinger, 2017, USA). Pyronaridine and studied analogs were docked at all sites using Glide in extra precision mode (XP mode) with default parameters (*73*). The site with the highest *gscore* was considered the binding site for each compound. A similar protocol was also performed using Haddock v2.4 (*74*). Electrostatic potential was calculated with APBS. Figures were generated with ChimeraX.

### Molecule synthesis

**Scheme 1.**
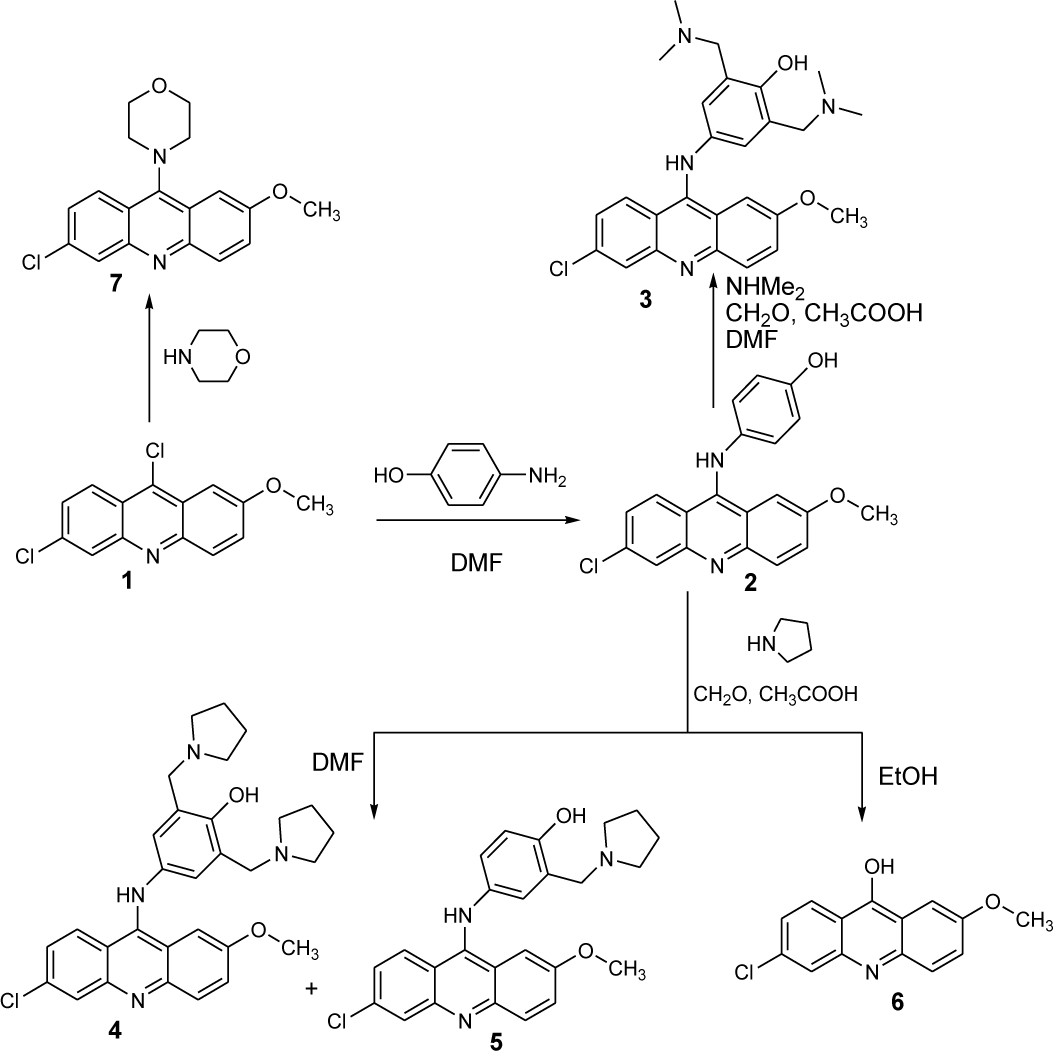
Synthetic route for 6-chloro-2-methoxyacridine derivatives

**Scheme 2.**
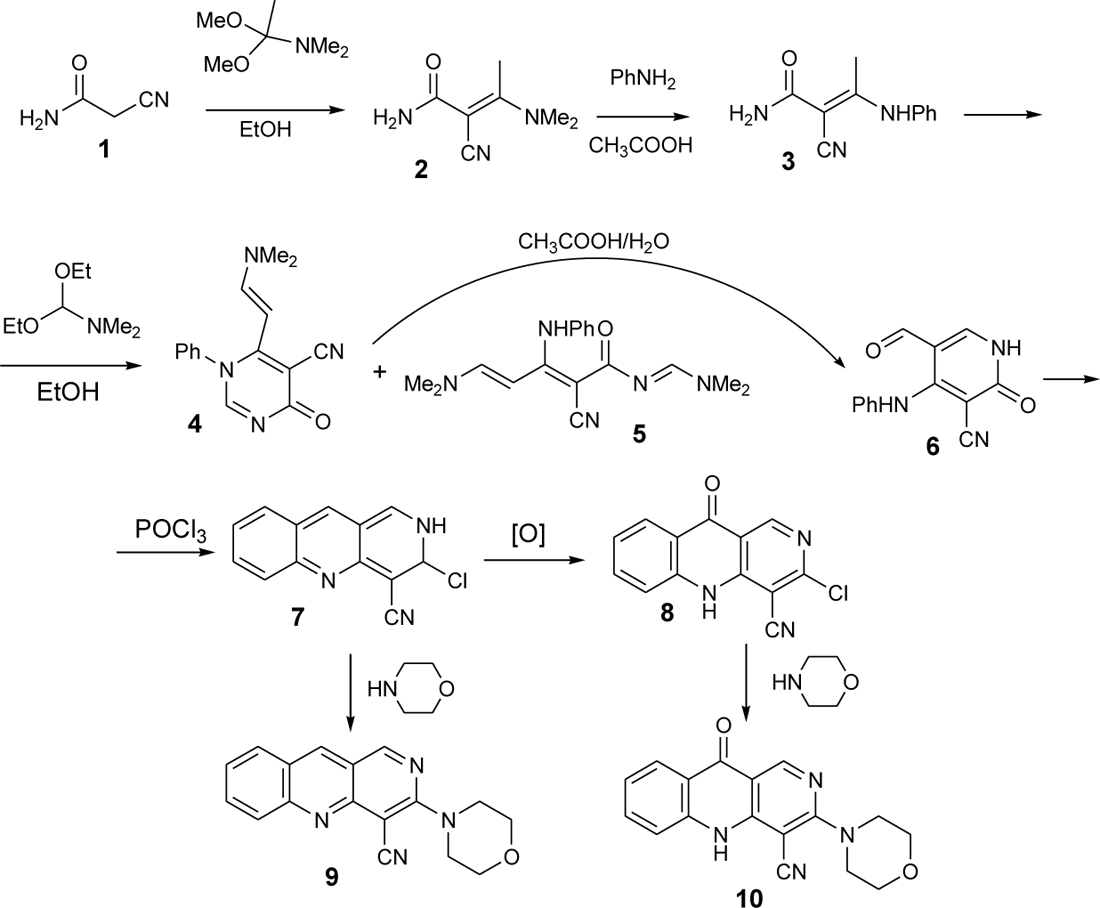
Synthetic route for benzo[b]-1,6-naphthyridine derivatives

### 4-[(6-chloro-2-methoxyacridin-9-yl)amino]phenol 2. (12126037)

Solid 2-aminophenol (0.39 g, 3.6 mmol) was added to a suspension of starting 6,9-dichloro-2-methoxyacridine 1 (0.5 g, 1.8 mmol) in 10 ml of dimethylformamide. The reaction mass was stirred at reflux for 1 hour. Cooled, the precipitate was filtered off, wash with DMF, EtOH and diethyl ether. The yield 72%. Mp 290 °C with decomposition, (DMF). Mass (EI), m / z (Irelat. (%)): 350.7981 [M]^+^ (92). C_20_H_15_ClN_2_O_2_. ^1^H NMR (DMSO-d_6_; δ, ppm): 9.98 (1H, s, NH), 8.16 (3H, m, 3CH), 7.73 (2H, m, 2CH), 7.47 (1H, d, *J* = 4.5, CH), 7.23 (2H, d, *J* = 6.5, 2CH), 6.92 (2H, d, *J* = 6.5, 2CH), 3.81 (3H, s, OCH_3_). ^13^C NMR(DMSO-d_6_; δ, ppm): 156.0, 152.9, 147.8, 147.6, 145.1, 144.7, 133.7, 131.3, 131.1, 128.1,122.8, 121.8, 121.5, 121.4, 120.9, 120.3, 120.1, 117.9, 101.0, 55.7.

### 4-[(6-chloro-2-methoxyacridin-9-yl)amino]-2,6-bis[(dimethylamino)methyl]phenol**3**. (12126038)

A mixture of 40% aqueous solution of dimethylamine (1.1 ml, 9.0 mmol), 1 ml of acetic acid and 37% aqueous solution of formaldehyde (0.39 ml, 5.4 mmol) were added at once to a suspension of 4-[(6-chloro-2-methoxyacridin-9-yl)amino]phenol 2 (0.9 mmol) in 10 ml of dimethylformamide. The suspension was heated to boiling and boiled for 4 hours. The reaction mixture was cooled, an aqueous solution of NaHCO_3_ (17 mmol) is added, transferred to a separatory funnel and extracted with ethyl acetate (30 ml x 2). Combined organic layer was washed with water 2 × 100 ml and dried over sodium sulfate. The solvent was removed in vacuo and residue oil was treated by mixture of 20 ml of hexane and 1 ml of diethyl ether. The oil was ground and kept for 10 hours at 40 °C. The solid was filtered off, washed with hexane and purified by column chromatography (methanol). The yield of aim product was 25%. Mp 148-152 °С. Mass (EI), *m/z* (*I_relat_.*(%)): 464.9869 [M]^+^ (88). C_26_H_29_ClN_4_O_2_. ^1^H NMR (DMSO-d_6_; δ, ppm): 9.98 (1H, s, NH), 8.17 (3H, m, 3CH), 7.84 (2H, m, 2CH), 7.45 (1H, d, *J* = 4.5, CH), 7.06 (2H, s, 2CH), 3.81 (3H, s, OCH_3_), 3.61 (2H, s, 2CH_2_), 2.78 (12H, br s, 2N(CH_3_)_2_. ^13^C NMR (DMSO-d_6_; δ, ppm): 157.2, 156.0, 154.5, 147.8, 145.1, 144.6, 133.7, 131.3, 131.1, 128.0, 128.1, 125.5, 122.8, 121.4, 118.6, 114.9, 112.2, 101.0, 55.8, 55.7, 44.9.

### 4-[(6-chloro-2-methoxyacridin-9-yl)amino]-2-(pyrrolidin-1-ylmethyl)phenol 4 ***and*** 4-***[(6-chloro-2-methoxyacridin-9-yl)amino]-2,6-bis(pyrrolidin-1-ylmethyl)phenol 5*.**

A mixture of pyrrolidine (0.52 ml, 6.0 mmol), 0.7 ml of acetic acid, and 37% aqueous formaldehyde solution (0.28 ml, 3.6 mmol) were added to a suspension of 4-[(6-chloro-2-methoxyacridin-9-yl)amino]phenol 2 (0.6 mmol) in 10 ml of dimethylformamide. The solution was stirred at reflux for 1 hour. The reaction mixture was cooled, an aqueous solution of NaHCO_3_ (11 mmol) was added and extracted with ethyl acetate (2 x 30 ml). The combined organic layer was washed with water (2 × 100 ml), dried by sodium sulfate and solvent was evaporated in vacuo. The resulting oil was dissolved in chloroform and applied to a chromatography column (chloroform/methanol 9:1) and product 4 was isolated. Pure methanol was used as eluent for separation product 5.

### 4-[(6-chloro-2-methoxyacridin-9-yl)amino]-2-(pyrrolidin-1-ylmethyl)phenol 4. (12126040)

The yield is 5%. Mp 178-182 °С. Mass (EI), *m/z* (*I_relat_.*(%)): 433.9298 [M]^+^ (76). C_25_H_24_ClN_3_O_2_. ^1^H NMR (DMSO-d_6_; δ, ppm): 10.21 (1H, s, NH), 8.17 (3H, m, 3CH), 7.81 (2H, m, 2CH), 7.45 (1H, d, *J* = 4.5, CH), 7.06 (2H, s, 2CH), 3.83 (3H, s, OCH_3_), 3.60 (2H, s, 2CH_2_), 2.5-2.7 (8H, br m, 2N(CH_2_)_2_), 1.6-1.8 (8H, br m, 2(CH_2_)_2_). ^13^C NMR (DMSO-d_6_; δ, ppm): 157.7, 156.2, 154.5, 147.8, 145.1, 144.3, 133.7, 131.3, 131.1, 125.5, 122.8, 121.4, 118.7, 114.9, 111.5, 101.1, 55.7, 54.1, 52.3, 23.4.

### 4-[(6-chloro-2-methoxyacridin-9-yl)amino]-2,6-bis(pyrrolidin-1-ylmethyl)phenol 5. (12126039)

The yield is 10%. Mp 145-150 °С. Mass (EI), *m/z* (*I_relat_.*(%)): 517.0615 [M]^+^ (79). C_30_H_33_ClN_4_O_2_. ^1^H NMR (DMSO-d_6_; δ, ppm): 10.04 (1H, s, NH), 8.16 (3H, m, 3CH), 7.79 (2H, m, 2CH), 7.47 (1H, d, *J* = 4.5, CH), 7.06 (1H, d, *J* = 4.5, CH), 6.93 (1H, s, CH), 3.83 (3H, s, OCH_3_), 3.60 (2H, s, 2CH_2_), 2.5-2.7 (8H, br m, 2N(CH_2_)_2_), 1.6-1.8 (8H, br m, 2(CH_2_)_2_). ^13^C NMR (DMSO-d_6_; δ, ppm): 156.0, 152.4, 149.3, 147.7, 145.1, 139.2, 131.2, 131.1, 128.7, 124.1, 122.8, 121.2, 120.8, 116.4, 116.1, 112.4, 101.6, 55.8, 51.9, 23.42.

### 6-chloro-2-methoxyacridin-9-ol 6. (12126072)

The mixture of pyrrolidine (1.0 ml, 13.0 mmol), 1.4 ml of acetic acid and 37% aqueous formaldehyde solution (0.56 ml, 7.8 mmol) were added to a suspension of 4-[(6-chloro-2-methoxyacridin-9-yl)amino]phenol 2 (1.3 mmol) in 10 ml of ethanol. The suspension was boiled for 36 hours. The reaction mixture was cooled, an aqueous solution of NaHCO_3_ (2.6 mmol) was added and the precipitate of aim compound was filtered off. The yield is 38%. Mp 300 °С (DMF). Mass (EI), *m/z* (*I_relat_.*(%)): 259.0615 [M]^+^ (79). C_14_H_10_ClNO_2_. ^1^H NMR (DMSO-d_6_; δ, ppm): 8.56 (1H, d, *J* = 9.0, CH), 7.72 (H, s, CH), 7.62 (1H, d, *J* = 4.5, CH), 7.44 (1H, d, *J* = 4.5, CH), 7.27 (1H, s, CH), 7.07 (1H, d, *J* = 9.0, CH), 3.86 (3H, s, OCH_3_). ^13^C NMR (DMSO-d_6_; δ, ppm): 157.8, 148.2, 147.9, 140.6, 138.0, 136.7, 128.7, 126.9, 125.5, 124.6, 123.3, 102.9, 55.3.

### 6-chloro-2-methoxy-9-morpholin-4-ylacridine 7. (12126036)

Morpholine (0.23 ml, 2.7 mmol) was added to a suspension of starting 6,9-dichloro-2-methoxyacridine 1 (0.25 g, 0.9 mmol) in 7 ml of dimethylformamide. The reaction mixture was stirred at reflux for 30 min, cooled, formed precipitate was filtered off and washed with dimethylformamide, acetone. The yield is 70%. Mp 208-212 °С (DMF). Mass (EI), *m/z* (*I_relat_.*(%)): 328.7926 [M]^+^ (74). C_18_H_17_ClN_2_O_2_. ^1^H NMR (DMSO-d_6_; δ, ppm): 8.56 (1H, d, *J* = 9.0, CH), 7.72 (H, s, CH), 7.62 (1H, d, *J* = 4.5, CH), 7.44 (1H, d, *J* = 4.5, CH), 7.27 (1H, s, CH), 7.07 (1H, d, *J* = 9.0, CH), 3.86 (3H, s, OCH_3_). ^13^C NMR (DMSO-d_6_; δ, ppm):155.7, 149.8, 142.6, 134.5, 131.7, 130.9, 130.6, 127.6, 122.4, 119.7, 119.1, 114.8, 100.5, 66.8, 55.7, 50.8.

### (2E)-2-cyano-3-(dimethylamino)but-2-enamide *2*

Dimethyl acetal dimethylacetamide (1 ml, 7.2 mmol) was added to a suspension of cyanoacetamide 1 (0.5 g, 6 mmol) in 15 ml of absolute alcohol. The suspension was boiled for 3 hours, cooled, obtained precipitate was filtered off, washed with alcohol and diethyl ether. Aim product 2 was obtained with a yield of 82%. Mp 188-192 °С (isopropanol).

### (2E)-3-anilino-2-cyanobut-2-enamide *3*

Aniline (0.45 ml, 5 mmol) was added to a suspension of (2E)-2-cyano-3-(dimethylamino)but-2-enamide 2 (0.3 g, 2 mmol) in acetic acid (4 ml). The suspension was boiled for 3 hours, cooled, acetic acid was removed under vacuum/ The residue was treated by water, formed precipitate was filtered off and washed with water, isopropyl alcohol and diethyl ether. Aim product 3 was obtained with a yield of 87%. Mp 179-182 °С (isopropanol).

### *A mixture of* 6-[(E)-2-(dimethylamino)vinyl]-4-oxo-1-phenyl-1,4-dihydropyrimidine-5-carbonitrile *4* ***and*** (2Z,4E)-3-anilino-2-cyano-5-(dimethylamino)-N-[(1E)-(dimethylamino)methylene]penta-2,4-dienamide *5*

Dimethylformamide diethyl acetal (1.3 ml, 7.5 mmol) was added to a suspension of (2E)-3-anilino-2-cyanobut-2-enamide 3 (0.6 g, 3 mmol) in 5 ml of absolute ethanol. The dark red solution was boiled for 6 hours. Part of the solvent was removed under vacuum until a thick suspension was obtained. The precipitate was filtered off, washed with absolute alcohol and dry diethyl ether. A mixture of compounds 4 and 5 was obtained.

### 4-anilino-5-formyl-2-oxo-1,2-dihydropyridine-3-carbonitrile *6*

A mixture of 6-[(E)-2-(dimethylamino)vinyl]-4-oxo-1-phenyl-1,4-dihydropyrimidine-5-carbonitrile 4 and (2Z,4E)-3-anilino-2-cyano-5-(dimethylamino)-N-[(1E)-(dimethylamino)methylene]penta-2,4-dienamide 5 (5.6 g) is dissolved in 56 ml of 90% acetic acid. A precipitate form after 30 min and reaction mixture was stirred at room temperature for 20 h. The precipitate was filtered off, washed with water, ethyl alcohol and acetone. The yield is 2.3 g of aim product 6. Mp 225-230 °C (with decomposition).

### 3-chloro-2,3-dihydrobenzo[b]-1,6-naphthyridine-4-carbonitrile *7*

The 4-anilino-5-formyl-2-oxo-1,2-dihydropyridine-3-carbonitrile 6 (2.3 g) was refluxed in phosphorus oxychloride (22 ml) for 1 hour. The reaction mixture was poured onto ice, stirred for 30 min. Formed precipitate was filtered off, washed with water, ethyl alcohol, diethyl ether. Aim product 7 was obtained in 80% yield. Mp 300-304 °С (DMF).

### 3-chloro-10-oxo-5,10-dihydrobenzo[b]-1,6-naphthyridine-4-carbonitrile *8*

To a suspension of 3-chloro-2,3-dihydrobenzo[b]-1,6-naphthyridine-4-carbonitrile 7 (0.44g) in 15 ml of acetone was added m-chloroperbenzoic acid 55% (1.45 g) in small portions and refluxed for 9 hours. The reaction mixture was cooled, obtained precipitate was filtered off and washed with acetone, toluene, acetone. The aim product 8 was obtained with a yield 50%. Mp 305 °C.

### 3-morpholin-4-ylbenzo[b]-1,6-naphthyridine-4-carbonitrile 9. (12126035)

Morpholine (0.16 ml, 1.8 mmol) was added to a suspension of 3-chloro-2,3-dihydrobenzo[b]-1,6-naphthyridine-4-carbonitrile 7 (0.15 g, 0.6 mmol) in 4 ml of dry dimethylformamide. The solution was boiled for 1 hour. Cooled, poured into 40 ml of water, extracted 2 times with 30 ml of ethyl acetate. The organic fractions were additionally washed with water 2 times. Ethyl acetate was removed under vacuum, ethyl alcohol was added to the dry residue, and the precipitate was filtered off. Washed with diethyl ether. The aim product 9 was obtained with a yield of 78%.Mp 222-225 °С (EtOH:DMF 2:1). Mass (EI), *m/z* (*I_relat_.*(%)): 290.3194 [M]^+^ (79). C_17_H_14_N_4_O. ^1^H NMR (DMSO-d_6_; δ, ppm): 9.46 (1H, s, CH), 9.13 (1H, s, CH), 8.16 (1H, d, *J* = 6.5, CH), 7.91 (1H, s, CH), 7.44 (1H, d, *J* = 6.5, CH), 4.16 (4H, m, O(CH_2_)_2_), 3.78 (4H, m, N(CH_2_)_2_). ^13^C NMR (DMSO-d_6_; δ, ppm): 144.9, 143.5, 137.8, 136.4, 136.0, 133.1, 129.5, 128.4, 126.8, 126.1, 114.5, 101.4, 66.7, 53.8.

### 3-morpholin-4-yl-10-oxo-5,10-dihydrobenzo[b]-1,6-naphthyridine-4-carbonitrile *10*. (**10326099**)

Morpholine (0.14 ml, 1.6 mmol) was added to a solution of 3-chloro-10-oxo-5,10-dihydrobenzo[b]-1,6-naphthyridine-4-carbonitrile 8 (0.2 g, 0.8 mmol) in 5 ml of dry dimethylformamide. Formed suspension was stirred at reflux for 1 hour, cooled, the precipitate was filtered off, washed with dimethylformamide and acetone. Mp 290 °С (из DMF). The yield is 38%. Mass (EI), *m/z* (*I_relat_.*(%)): 306.3188 [M]^+^ (68). C_17_H_14_N_4_O_2_. ^1^H NMR (DMSO-d_6_; δ, ppm): 8.02 (1H, s, CH), 7.91 (1H, d, *J* = 6.5, CH), 7.59 (1H, s, CH),7.09 (1H, d, *J* = 6.5, CH), 4.08 (4H, m, O(CH_2_)_2_), 3.72 (4H, m, N(CH_2_)_2_). ^13^C NMR (DMSO-d_6_; δ, ppm): 182.5, 146.4, 146.1, 139.7, 139.1, 138.7, 137.0, 124.9, 122.7, 118.3, 115.4,108.8, 94.9, 66.8, 53.7.

### PL^pro^ cloning, expression, purification and assays

The viral cDNA template (GenBank MT126808.1) was kindly provided by Dr. Edison Durigon. Amplification of the nucleotide sequence coding for the PL^pro^ domain (residues 1564-1879 of SARSCoV-2) was performed by Polymerase Chain Reaction (PCR) using forward (5’-ATTCCATGGGCGAAGTGAGGACTATTAAGGTGTTTAC-3’) and reverse (5’-ATTGCTCGAGTGGTTTTATGGTTGTTGTGTAACT-3’) primers, with restriction sites for *NcoI* and *XhoI*. PCR product was digested with *NcoI* and *XhoI* and cloned into pET28a (Novagen) in frame with a C-terminal his-tag coding sequence *E. coli* BL21 transformed with plasmids were grown in LB to an optical density (OD_600_) of 0.6 at 37 °C and 200 RPM. Protein expression was induced adding 0.5 mM IPTG and 1 mM Zinc Chloride and grown overnight at 18°. The cell pellet was resuspended in lysis buffer (50 mM Tris, 150 mM NaCl, 10 mM imidazole, 1 mM DTT, pH 8.5), disrupted by sonication and centrifuged at 12,500 rpm for 40 min at 4° C. Protein was isolated from the lysate using 5 mL Ni-NTA resin (Qiagen) pre-equilibrated with lysis buffer and then washed with 15 column volumes (CV) with the same buffer. The his-tagged protein was eluted 4 CV of elution buffer (lysis buffer supplemented with 250 mM imidazole) and further purified by size exclusion chromatography in a Superdex 200 10/30 (GE Healthcare) equilibrated with 20 mM Tris pH 7.4, 100 mM NaCl, 2 mM DTT.

PL^pro^ inhibition assay was performed using the FRET-based fluorescent peptide substrate Abz-TLKGG↓APIKEDDPS-EDDnp, kindly provided by Dr. Maria Aparecida Juliano (Federal University of São Paulo, Brazil). The assay was standardized with enzyme concentration of 70 nM and 27 µM of fluorescent substrate in PL^pro^ assay buffer (50 mM HEPES pH 7.5, 0.01 % Triton X-100 and 5 mM DTT), at 37°C for 30 min. Activity was measured in the plate reader system Spectramax Gemini EM (Molecular devices), with 11_ex_= 320 nm and 11_em_= 420 nm, in presence of different inhibitors and 1% DMSO.

### M^pro^ cloning, expression, purification and assays

The M^pro^ cloning is described elsewhere (*75*). For expression, BL21 cells transformed with plasmid were grown in ZYM-5052 to an OD_600_ of 0.6-0.8 at 37°C and 200 RPM. Protein expression was induced lowering the temperature to 18 °C, cells were then grown for 16h and harvested by centrifugation. The cell pellet was resuspended in lysis buffer (20 mM Tris pH 7.8, 150 mM NaCl, 1 mM DTT), disrupted by sonication and centrifuged at 12,000 x g for 40 min at 4 °C. Uncleaved protein with 6xHisTag were isolated from the lysate using Ni-NTA resin (Qiagen). The flow-through (FT) was then used to purify cleaved protein with ammonium sulfate precipitation. Ammonium sulfate was added into the FT to a final concentration of 1M, incubated on ice for 10 min and centrifuged at 12,000 x g for 30 min at 4 °C. The precipitated protein was resuspended in gel filtration buffer (20 mM Tris pH 7.8, 150 mM NaCl, 1 mM EDTA, 1 mM DTT) and purified by size-exclusion chromatography in a HiLoad 26/100 Superdex 200 column (GE Healthcare) pre-equilibrated with gel filtration buffer. Afterwards, the protein was buffer exchanged to 20 mM Tris pH 8.0, 1.0 mM DTT and loaded into a Mono-Q 5/50 GL column (GE Healthcare) in the same buffer, and eluted with a linear gradient of a buffer containing 20 mM Tris pH 8.0, 1.0 M NaCl and 1.0 mM DTT. Proteins were concentrated at 1.0 mg/mL, flash-frozen and stored at −80°C for inhibition assays.

M^pro^ inhibition assay was performed using the FRET-based fluorescent peptide substrate DABCYL-KTSAVLQ↓SGFRKM-E(EDANS)-NH2 (purchased from Genscript). The assay was standardized with enzyme concentration of 140 nM and 30 µM of fluorescent substrate in M^pro^ assay buffer containing (20 mM Tris pH 7.3, 1 mM EDTA, 1 mM DTT), at 37°C for 30 min. Activity was detected in the spectrofluorometer system Spectramax Gemini EM (Molecular devices), with 11_ex_= 360 nm and 11_em_ 460 nm, in presence of different inhibitors and 1% DMSO.

### Kinase profiling of pyronaridine

Kinase profiling was performed for pyronaridine (1 μM) in duplicate by ThermoFisher (Life Technologies Corporation, Chicago, IL 60693) using Z’Lyte (*76*), Adapta (*77*) and LanthaScreen (*78*) assays for 485 purified kinases.

### CAMK1 inhibition

Pyronaridine inhibition of CAMK1 was performed by Selected Services (Thermo Fisher) using the the Adapta universal kinase assay, which is a homogenous, fluorescent-based immunoassay for the detection of ADP. The 2X CAMK1 (CaMK1) and ZIPtide mixture was prepared in 50 mM HEPES pH 7.5, 0.01% BRIJ-35, 10 mM MgCl2, 1 mM EGTA, 4 mM CaCl2, 800 U/ml Calmodulin, 0.02% NaN3. The final 10 μL Kinase Reaction consists of 0.25 - 1.2 ng CAMK1 (CaMK1) and 200 μM ZIPtide in 32.5 mM HEPES pH 7.5, 0.005% BRIJ-35, 5 mM MgCl2, 500 μM EGTA, 2 mM CaCl2, 400 U/ml Calmodulin, 0.01% NaN3. After the 1 h kinase reaction incubation, 5 μL of Detection Mix is added.

## ABBREVIATIONS USED

ACE2: Angiotensin converting enzyme 2

BMP: Bis(monoacylglycero)phosphate

COVID-19: Coronavirus disease

MERS-CoV: Middle East Respiratory Syndrome coronavirus

SARS-CoV-2: Severe Acute Respiratory coronavirus 2

## Supplemental Material

**Table S1.** SelectScreen kinase profiling of pyronaridine at 1µM tested in duplicate.

| **Kinase** | **% inhibition**<br>**mean** | **assay** |
| --- | --- | --- |
| CAMK1 (CaMK1) | 35 | Adapta |
| CDK4/cyclin D1 | 1 | Adapta |
| CDK4/cyclin D3 | -6 | Adapta |
| CDK6/cyclin D1 | 1 | Adapta |
| CDK7/cyclin H/MNAT1 | -13 | Adapta |
| CDK9/cyclin T1 | 5 | Adapta |
| CHUK (IKK alpha) | 0 | Adapta |
| DAPK1 | 11 | Adapta |
| GSG2 (Haspin) | 9 | Adapta |
| IRAK1 | -4 | Adapta |
| LRRK2 FL | -4 | Adapta |
| LRRK2 G2019S FL | -10 | Adapta |
| LRRK2 G2019S | 4 | Adapta |
| LRRK2 I2020T | -15 | Adapta |
| LRRK2 R1441C | -7 | Adapta |
| LRRK2 | -2 | Adapta |
| NUAK1 (ARK5) | 11 | Adapta |
| PI4K2A (PI4K2 alpha) | 4 | Adapta |
| PI4K2B (PI4K2 beta) | 2 | Adapta |
| PI4KA (PI4K alpha) | 0 | Adapta |
| PI4KB (PI4K beta) | 5 | Adapta |
| PIK3C2A (PI3K-C2 alpha) | -1 | Adapta |
| PIK3C2B (PI3K-C2 beta) | -5 | Adapta |
| PIK3C2G (PI3K-C2 gamma) | 3 | Adapta |
| PIK3C3 (hVPS34) | -4 | Adapta |
| PIK3CA E542K/PIK3R1 (p110 alpha<br>E542K/p85 alpha) | -4 | Adapta |
| PIK3CA E545K/PIK3R1 (p110 alpha<br>E545K/p85 alpha) | 3 | Adapta |
| PIK3CA/PIK3R1 (p110 alpha/p85 alpha) | 7 | Adapta |
| PIK3CA/PIK3R3 (p110 alpha/p55 gamma) | 1 | Adapta |
| PIK3CB/PIK3R1 (p110 beta/p85 alpha) | -11 | Adapta |
| PIK3CB/PIK3R2 (p110 beta/p85 beta) | -2 | Adapta |
| PIK3CD/PIK3R1 (p110 delta/p85 alpha) | 9 | Adapta |
| PIK3CG (p110 gamma) | 2 | Adapta |
| PIP4K2A | 15 | Adapta |
| PIP5K1A | -6 | Adapta |
| PIP5K1B | -11 | Adapta |
| PIP5K1C | -3 | Adapta |
| SPHK1 | 10 | Adapta |
| SPHK2 | -5 | Adapta |
| AAK1 | 0 | Lanthascreen |
| ABL1 H396P | 7 | Lanthascreen |
| ABL1 M351T | 8 | Lanthascreen |
| ABL1 Q252H | -10 | Lanthascreen |
| ACVR1 (ALK2) R206H | 2 | Lanthascreen |
| ACVR1 (ALK2) | 0 | Lanthascreen |
| ACVR2A | -6 | Lanthascreen |
| ACVR2B | 8 | Lanthascreen |
| ACVRL1 (ALK1) | 12 | Lanthascreen |
| ADCK3 | -16 | Lanthascreen |
| ALK C1156Y | -2 | Lanthascreen |
| ALK F1174L | 8 | Lanthascreen |
| ALK L1196M | -3 | Lanthascreen |
| ALK R1275Q | 6 | Lanthascreen |
| ALK T1151_L1152insT | 4 | Lanthascreen |
| AMPK (A1/B1/G2) | -7 | Lanthascreen |
| AMPK (A1/B1/G3) | -1 | Lanthascreen |
| AMPK (A1/B2/G1) | -2 | Lanthascreen |
| AMPK (A2/B2/G1) | 6 | Lanthascreen |
| AMPK (A2/B2/G2) | 3 | Lanthascreen |
| ANKK1 | -2 | Lanthascreen |
| AXL R499C | 3 | Lanthascreen |
| BMPR1A (ALK3) | 10 | Lanthascreen |
| BMPR1B (ALK6) | -3 | Lanthascreen |
| BMPR2 | 8 | Lanthascreen |
| BRAF V599E | 4 | Lanthascreen |
| BRAF | -11 | Lanthascreen |
| BRSK2 | 4 | Lanthascreen |
| CAMK2G (CaMKII gamma) | -1 | Lanthascreen |
| CAMKK1 (CAMKKA) | 15 | Lanthascreen |
| CAMKK2 (CaMKK beta) | -1 | Lanthascreen |
| CASK | 2 | Lanthascreen |
| CDC7/DBF4 | -6 | Lanthascreen |
| CDK11 (Inactive) | -1 | Lanthascreen |
| CDK11/cyclin C | -18 | Lanthascreen |
| CDK13/cyclin K | -13 | Lanthascreen |
| CDK14 (PFTK1)/cyclin Y | 14 | Lanthascreen |
| CDK16 (PCTK1)/cyclin Y | -12 | Lanthascreen |
| CDK2/cyclin A1 | 5 | Lanthascreen |
| CDK2/cyclin E1 | 0 | Lanthascreen |
| CDK2/cyclin O | 2 | Lanthascreen |
| CDK3/cyclin E1 | 3 | Lanthascreen |
| CDK5 (Inactive) | -1 | Lanthascreen |
| CDK8/cyclin C | 3 | Lanthascreen |
| CDK9 (Inactive) | -10 | Lanthascreen |
| CDK9/cyclin K | 5 | Lanthascreen |
| CLK4 | 6 | Lanthascreen |
| DAPK2 | -6 | Lanthascreen |
| DDR1 | -2 | Lanthascreen |
| DDR2 N456S | 17 | Lanthascreen |
| DDR2 T654M | -11 | Lanthascreen |
| DDR2 | 0 | Lanthascreen |
| DMPK | 4 | Lanthascreen |
| DYRK2 | -7 | Lanthascreen |
| EGFR (ErbB1) d746-750 | -10 | Lanthascreen |
| EGFR (ErbB1) d747-749 A750P | 0 | Lanthascreen |
| EIF2AK2 (PKR) | 4 | Lanthascreen |
| EPHA3 | 11 | Lanthascreen |
| EPHA6 | -3 | Lanthascreen |
| EPHA7 | -3 | Lanthascreen |
| ERN1 | 6 | Lanthascreen |
| ERN2 | 6 | Lanthascreen |
| FGFR1 V561M | 5 | Lanthascreen |
| FGFR3 G697C | 7 | Lanthascreen |
| FGFR3 K650M | -4 | Lanthascreen |
| FLT3 ITD | 9 | Lanthascreen |
| FYN A | -3 | Lanthascreen |
| GAK | 28 | Lanthascreen |
| GRK1 | 1 | Lanthascreen |
| HUNK | -1 | Lanthascreen |
| ICK | 0 | Lanthascreen |
| IRAK3 | -5 | Lanthascreen |
| KIT A829P | -9 | Lanthascreen |
| KIT D816H | -7 | Lanthascreen |
| KIT D816V | -3 | Lanthascreen |
| KIT D820E | -4 | Lanthascreen |
| KIT N822K | 14 | Lanthascreen |
| KIT T670E | 1 | Lanthascreen |
| KIT V559D T670I | -10 | Lanthascreen |
| KIT V654A | 10 | Lanthascreen |
| KIT Y823D | -4 | Lanthascreen |
| LATS2 | -4 | Lanthascreen |
| LIMK1 | -7 | Lanthascreen |
| LIMK2 | -1 | Lanthascreen |
| MAP2K1 (MEK1) S218D S222D | 3 | Lanthascreen |
| MAP2K1 (MEK1) | 1 | Lanthascreen |
| MAP2K2 (MEK2) | -3 | Lanthascreen |
| MAP2K4 (MEK4) | 1 | Lanthascreen |
| MAP2K5 (MEK5) | 1 | Lanthascreen |
| MAP2K6 (MKK6) S207E T211E | 10 | Lanthascreen |
| MAP2K6 (MKK6) | 9 | Lanthascreen |
| MAP3K10 (MLK2) | 0 | Lanthascreen |
| MAP3K11 (MLK3) | 2 | Lanthascreen |
| MAP3K14 (NIK) | 2 | Lanthascreen |
| MAP3K2 (MEKK2) | 7 | Lanthascreen |
| MAP3K3 (MEKK3) | -3 | Lanthascreen |
| MAP3K5 (ASK1) | -11 | Lanthascreen |
| MAP3K7/MAP3K7IP1 (TAK1-TAB1) | 1 | Lanthascreen |
| MAP4K1 (HPK1) | 1 | Lanthascreen |
| MAP4K3 (GLK) | 1 | Lanthascreen |
| MAPK10 (JNK3) | 5 | Lanthascreen |
| MAPK15 (ERK7) | -11 | Lanthascreen |
| MAPK8 (JNK1) | 3 | Lanthascreen |
| MAPK9 (JNK2) | 6 | Lanthascreen |
| MASTL | 3 | Lanthascreen |
| MERTK (cMER) A708S | 0 | Lanthascreen |
| MET D1228H | 3 | Lanthascreen |
| MKNK2 (MNK2) | 2 | Lanthascreen |
| MLCK (MLCK2) | -6 | Lanthascreen |
| MLK4 | 8 | Lanthascreen |
| MYLK (MLCK) | 4 | Lanthascreen |
| MYLK4 | -9 | Lanthascreen |
| MYO3A (MYO3 alpha) | -6 | Lanthascreen |
| MYO3B (MYO3 beta) | 5 | Lanthascreen |
| NEK8 | 0 | Lanthascreen |
| NLK | 8 | Lanthascreen |
| NUAK2 | 2 | Lanthascreen |
| PKMYT1 | 1 | Lanthascreen |
| PKN2 (PRK2) | 6 | Lanthascreen |
| PLK4 | 6 | Lanthascreen |
| PRKACB (PRKAC beta) | 12 | Lanthascreen |
| PRKACG (PRKAC gamma) | 2 | Lanthascreen |
| RAF1 (cRAF) Y340D Y341D | -5 | Lanthascreen |
| RET G691S | -3 | Lanthascreen |
| RET M918T | -3 | Lanthascreen |
| RET V804M | -1 | Lanthascreen |
| RIPK2 | 1 | Lanthascreen |
| RIPK3 | -7 | Lanthascreen |
| SIK1 | 2 | Lanthascreen |
| SIK3 | -2 | Lanthascreen |
| SLK | -1 | Lanthascreen |
| STK16 (PKL12) | 5 | Lanthascreen |
| STK17A (DRAK1) | 1 | Lanthascreen |
| STK17B (DRAK2) | -4 | Lanthascreen |
| STK32B (YANK2) | 0 | Lanthascreen |
| STK32C (YANK3) | 1 | Lanthascreen |
| STK33 | 11 | Lanthascreen |
| STK38 (NDR) | 2 | Lanthascreen |
| STK38L (NDR2) | -7 | Lanthascreen |
| STK39 (STLK3) | -19 | Lanthascreen |
| TAOK1 | -4 | Lanthascreen |
| TAOK3 (JIK) | -2 | Lanthascreen |
| TEC | 7 | Lanthascreen |
| TEK (TIE2) R849W | -3 | Lanthascreen |
| TEK (TIE2) Y1108F | -2 | Lanthascreen |
| TESK1 | -2 | Lanthascreen |
| TESK2 | -3 | Lanthascreen |
| TGFBR1 (ALK5) | -4 | Lanthascreen |
| TGFBR2 | -4 | Lanthascreen |
| TLK1 | 12 | Lanthascreen |
| TLK2 | -2 | Lanthascreen |
| TNIK | -1 | Lanthascreen |
| TNK2 (ACK) | 1 | Lanthascreen |
| TTK | 4 | Lanthascreen |
| ULK1 | -4 | Lanthascreen |
| ULK2 | 2 | Lanthascreen |
| ULK3 | 8 | Lanthascreen |
| VRK2 | -4 | Lanthascreen |
| WEE1 | 10 | Lanthascreen |
| WNK1 | 4 | Lanthascreen |
| WNK2 | 4 | Lanthascreen |
| WNK3 | 7 | Lanthascreen |
| ZAK | 0 | Lanthascreen |
| ABL1 E255K | 7 | Z'Lyte |
| ABL1 F317I | 0 | Z'Lyte |
| ABL1 F317L | -1 | Z'Lyte |
| ABL1 G250E | 1 | Z'Lyte |
| ABL1 T315I | 11 | Z'Lyte |
| ABL1 Y253F | -1 | Z'Lyte |
| ABL1 | 2 | Z'Lyte |
| ABL2 (Arg) | 2 | Z'Lyte |
| ACVR1B (ALK4) | -1 | Z'Lyte |
| ADRBK1 (GRK2) | -3 | Z'Lyte |
| ADRBK2 (GRK3) | -5 | Z'Lyte |
| AKT1 (PKB alpha) | 4 | Z'Lyte |
| AKT2 (PKB beta) | 1 | Z'Lyte |
| AKT3 (PKB gamma) | 7 | Z'Lyte |
| ALK | -1 | Z'Lyte |
| AMPK (A1/B2/G2) | -2 | Z'Lyte |
| AMPK (A1/B2/G3) | 4 | Z'Lyte |
| AMPK (A2/B1/G2) | 1 | Z'Lyte |
| AMPK (A2/B1/G3) | 2 | Z'Lyte |
| AMPK (A2/B2/G3) | 2 | Z'Lyte |
| AMPK A1/B1/G1 | 5 | Z'Lyte |
| AMPK A2/B1/G1 | 2 | Z'Lyte |
| AURKA (Aurora A) | 9 | Z'Lyte |
| AURKB (Aurora B) | 5 | Z'Lyte |
| AURKC (Aurora C) | -11 | Z'Lyte |
| AXL | 1 | Z'Lyte |
| BLK | 3 | Z'Lyte |
| BMX | 11 | Z'Lyte |
| BRAF V599E | 0 | Z'Lyte |
| BRAF | 4 | Z'Lyte |
| BRSK1 (SAD1) | 2 | Z'Lyte |
| BTK | 9 | Z'Lyte |
| CAMK1D (CaMKI delta) | -26 | Z'Lyte |
| CAMK1G (CAMKI gamma) | 0 | Z'Lyte |
| CAMK2A (CaMKII alpha) | 4 | Z'Lyte |
| CAMK2B (CaMKII beta) | 1 | Z'Lyte |
| CAMK2D (CaMKII delta) | -5 | Z'Lyte |
| CAMK4 (CaMKIV) | -5 | Z'Lyte |
| CDC42 BPA (MRCKA) | 2 | Z'Lyte |
| CDC42 BPB (MRCKB) | -3 | Z'Lyte |
| CDC42 BPG (MRCKG) | 1 | Z'Lyte |
| CDK1/cyclin B | 5 | Z'Lyte |
| CDK17/cyclin Y | -2 | Z'Lyte |
| CDK18/cyclin Y | 1 | Z'Lyte |
| CDK2/cyclin A | 0 | Z'Lyte |
| CDK5/p25 | 0 | Z'Lyte |
| CDK5/p35 | 3 | Z'Lyte |
| CDKL5 | 4 | Z'Lyte |
| CHEK1 (CHK1) | 8 | Z'Lyte |
| CHEK2 (CHK2) | 0 | Z'Lyte |
| CLK1 | -1 | Z'Lyte |
| CLK2 | 3 | Z'Lyte |
| CLK3 | -3 | Z'Lyte |
| CSF1R (FMS) | -3 | Z'Lyte |
| CSK | -3 | Z'Lyte |
| CSNK1A1 (CK1 alpha 1) | 5 | Z'Lyte |
| CSNK1A1L | 3 | Z'Lyte |
| CSNK1D (CK1 delta) | 5 | Z'Lyte |
| CSNK1E (CK1 epsilon) R178C | 1 | Z'Lyte |
| CSNK1E (CK1 epsilon) | 1 | Z'Lyte |
| CSNK1G1 (CK1 gamma 1) | 0 | Z'Lyte |
| CSNK1G2 (CK1 gamma 2) | 4 | Z'Lyte |
| CSNK1G3 (CK1 gamma 3) | 2 | Z'Lyte |
| CSNK2A1 (CK2 alpha 1) | 7 | Z'Lyte |
| CSNK2A2 (CK2 alpha 2) | 5 | Z'Lyte |
| DAPK3 (ZIPK) | 2 | Z'Lyte |
| DCAMKL1 (DCLK1) | 0 | Z'Lyte |
| DCAMKL2 (DCK2) | -1 | Z'Lyte |
| DNA-PK | 5 | Z'Lyte |
| DYRK1A | 3 | Z'Lyte |
| DYRK1B | 1 | Z'Lyte |
| DYRK3 | 1 | Z'Lyte |
| DYRK4 | 0 | Z'Lyte |
| EEF2K | 1 | Z'Lyte |
| EGFR (ErbB1) C797S | -7 | Z'Lyte |
| EGFR (ErbB1) G719C | -4 | Z'Lyte |
| EGFR (ErbB1) G719S | -5 | Z'Lyte |
| EGFR (ErbB1) L858R | 5 | Z'Lyte |
| EGFR (ErbB1) L861Q | -4 | Z'Lyte |
| EGFR (ErbB1) T790M C797S L858R | 2 | Z'Lyte |
| EGFR (ErbB1) T790M L858R | 5 | Z'Lyte |
| EGFR (ErbB1) T790M | 2 | Z'Lyte |
| EGFR (ErbB1) | -5 | Z'Lyte |
| EPHA1 | 1 | Z'Lyte |
| EPHA2 | 5 | Z'Lyte |
| EPHA4 | -2 | Z'Lyte |
| EPHA5 | -3 | Z'Lyte |
| EPHA8 | 1 | Z'Lyte |
| EPHB1 | 2 | Z'Lyte |
| EPHB2 | 4 | Z'Lyte |
| EPHB3 | -1 | Z'Lyte |
| EPHB4 | 7 | Z'Lyte |
| ERBB2 (HER2) | 2 | Z'Lyte |
| ERBB4 (HER4) | -2 | Z'Lyte |
| FER | -21 | Z'Lyte |
| FES (FPS) | -10 | Z'Lyte |
| FGFR1 | -5 | Z'Lyte |
| FGFR2 N549H | 2 | Z'Lyte |
| FGFR2 | -1 | Z'Lyte |
| FGFR3 K650E | -4 | Z'Lyte |
| FGFR3 V555M | -1 | Z'Lyte |
| FGFR3 | -2 | Z'Lyte |
| FGFR4 | -14 | Z'Lyte |
| FGR | 11 | Z'Lyte |
| FLT1 (VEGFR1) | -6 | Z'Lyte |
| FLT3 D835Y | 2 | Z'Lyte |
| FLT3 | 3 | Z'Lyte |
| FLT4 (VEGFR3) | 3 | Z'Lyte |
| FRAP1 (mTOR) | 1 | Z'Lyte |
| FRK (PTK5) | -2 | Z'Lyte |
| FYN | 2 | Z'Lyte |
| GRK4 | 25 | Z'Lyte |
| GRK5 | 10 | Z'Lyte |
| GRK6 | -1 | Z'Lyte |
| GRK7 | 4 | Z'Lyte |
| GSK3A (GSK3 alpha) | 4 | Z'Lyte |
| GSK3B (GSK3 beta) | 8 | Z'Lyte |
| HCK | 4 | Z'Lyte |
| HIPK1 (Myak) | 1 | Z'Lyte |
| HIPK2 | -1 | Z'Lyte |
| HIPK3 (YAK1) | -2 | Z'Lyte |
| HIPK4 | 5 | Z'Lyte |
| IGF1R | 6 | Z'Lyte |
| IKBKB (IKK beta) | 1 | Z'Lyte |
| IKBKE (IKK epsilon) | 2 | Z'Lyte |
| INSR | 10 | Z'Lyte |
| INSRR (IRR) | -1 | Z'Lyte |
| IRAK4 | -6 | Z'Lyte |
| ITK | 2 | Z'Lyte |
| JAK1 | 0 | Z'Lyte |
| JAK2 JH1 JH2 V617F | 0 | Z'Lyte |
| JAK2 JH1 JH2 | -3 | Z'Lyte |
| JAK2 | 4 | Z'Lyte |
| JAK3 | 1 | Z'Lyte |
| KDR (VEGFR2) | 7 | Z'Lyte |
| KIT T670I | 1 | Z'Lyte |
| KIT V559D V654A | -1 | Z'Lyte |
| KIT V559D | -1 | Z'Lyte |
| KIT V560G | -5 | Z'Lyte |
| KIT | -2 | Z'Lyte |
| KSR2 | 2 | Z'Lyte |
| LCK | 12 | Z'Lyte |
| LTK (TYK1) | -1 | Z'Lyte |
| LYN A | -2 | Z'Lyte |
| LYN B | -2 | Z'Lyte |
| MAP2K1 (MEK1) | -2 | Z'Lyte |
| MAP2K2 (MEK2) | 2 | Z'Lyte |
| MAP2K6 (MKK6) | 3 | Z'Lyte |
| MAP3K19 (YSK4) | 3 | Z'Lyte |
| MAP3K8 (COT) | -6 | Z'Lyte |
| MAP3K9 (MLK1) | 17 | Z'Lyte |
| MAP4K2 (GCK) | 5 | Z'Lyte |
| MAP4K4 (HGK) | 1 | Z'Lyte |
| MAP4K5 (KHS1) | 5 | Z'Lyte |
| MAPK1 (ERK2) | -1 | Z'Lyte |
| MAPK10 (JNK3) | 0 | Z'Lyte |
| MAPK11 (p38 beta) | -8 | Z'Lyte |
| MAPK12 (p38 gamma) | 7 | Z'Lyte |
| MAPK13 (p38 delta) | 5 | Z'Lyte |
| MAPK14 (p38 alpha) Direct | 3 | Z'Lyte |
| MAPK14 (p38 alpha) | 1 | Z'Lyte |
| MAPK3 (ERK1) | 6 | Z'Lyte |
| MAPK7 (ERK5) | 5 | Z'Lyte |
| MAPK8 (JNK1) | 3 | Z'Lyte |
| MAPK9 (JNK2) | -15 | Z'Lyte |
| MAPKAPK2 | 10 | Z'Lyte |
| MAPKAPK3 | 17 | Z'Lyte |
| MAPKAPK5 (PRAK) | 3 | Z'Lyte |
| MARK1 (MARK) | -10 | Z'Lyte |
| MARK2 | 6 | Z'Lyte |
| MARK3 | 2 | Z'Lyte |
| MARK4 | -4 | Z'Lyte |
| MATK (HYL) | -2 | Z'Lyte |
| MELK | 31 | Z'Lyte |
| MERTK (cMER) | 2 | Z'Lyte |
| MET (cMet) Y1235D | 4 | Z'Lyte |
| MET (cMet) | -1 | Z'Lyte |
| MET M1250T | 0 | Z'Lyte |
| MINK1 | 7 | Z'Lyte |
| MKNK1 (MNK1) | 4 | Z'Lyte |
| MST1R (RON) | 2 | Z'Lyte |
| MST4 | 1 | Z'Lyte |
| MUSK | -10 | Z'Lyte |
| MYLK2 (skMLCK) | 7 | Z'Lyte |
| NEK1 | -1 | Z'Lyte |
| NEK2 | -4 | Z'Lyte |
| NEK4 | 4 | Z'Lyte |
| NEK6 | -3 | Z'Lyte |
| NEK9 | 0 | Z'Lyte |
| NIM1K | -2 | Z'Lyte |
| NTRK1 (TRKA) | 19 | Z'Lyte |
| NTRK2 (TRKB) | 1 | Z'Lyte |
| NTRK3 (TRKC) | 2 | Z'Lyte |
| PAK1 | 6 | Z'Lyte |
| PAK2 (PAK65) | -1 | Z'Lyte |
| PAK3 | -7 | Z'Lyte |
| PAK4 | 3 | Z'Lyte |
| PAK6 | -2 | Z'Lyte |
| PAK7 (KIAA1264) | -5 | Z'Lyte |
| PASK | -4 | Z'Lyte |
| PDGFRA (PDGFR alpha) | 2 | Z'Lyte |
| PDGFRA D842V | -1 | Z'Lyte |
| PDGFRA T674I | 0 | Z'Lyte |
| PDGFRA V561D | 2 | Z'Lyte |
| PDGFRB (PDGFR beta) | 0 | Z'Lyte |
| PDK1 Direct | 7 | Z'Lyte |
| PDK1 | 0 | Z'Lyte |
| PEAK1 | 1 | Z'Lyte |
| PHKG1 | -1 | Z'Lyte |
| PHKG2 | -2 | Z'Lyte |
| PIM1 | 8 | Z'Lyte |
| PIM2 | -4 | Z'Lyte |
| PIM3 | 12 | Z'Lyte |
| PKN1 (PRK1) | 1 | Z'Lyte |
| PLK1 | 6 | Z'Lyte |
| PLK2 | 5 | Z'Lyte |
| PLK3 | 28 | Z'Lyte |
| PRKACA (PKA) | 1 | Z'Lyte |
| PRKCA (PKC alpha) | 13 | Z'Lyte |
| PRKCB1 (PKC beta I) | 2 | Z'Lyte |
| PRKCB2 (PKC beta II) | 15 | Z'Lyte |
| PRKCD (PKC delta) | 3 | Z'Lyte |
| PRKCE (PKC epsilon) | 6 | Z'Lyte |
| PRKCG (PKC gamma) | -4 | Z'Lyte |
| PRKCH (PKC eta) | -9 | Z'Lyte |
| PRKCI (PKC iota) | -10 | Z'Lyte |
| PRKCN (PKD3) | 8 | Z'Lyte |
| PRKCQ (PKC theta) | 8 | Z'Lyte |
| PRKCZ (PKC zeta) | -1 | Z'Lyte |
| PRKD1 (PKC mu) | -2 | Z'Lyte |
| PRKD2 (PKD2) | -4 | Z'Lyte |
| PRKG1 | -6 | Z'Lyte |
| PRKG2 (PKG2) | 1 | Z'Lyte |
| PRKX | 5 | Z'Lyte |
| PTK2 (FAK) | 2 | Z'Lyte |
| PTK2B (FAK2) | -1 | Z'Lyte |
| PTK6 (Brk) | -4 | Z'Lyte |
| RAF1 (cRAF) Y340D Y341D | -9 | Z'Lyte |
| RET A883F | -1 | Z'Lyte |
| RET S891A | -3 | Z'Lyte |
| RET V804E | 2 | Z'Lyte |
| RET V804L | 1 | Z'Lyte |
| RET Y791F | 5 | Z'Lyte |
| RET | 5 | Z'Lyte |
| ROCK1 | 2 | Z'Lyte |
| ROCK2 | 1 | Z'Lyte |
| ROS1 | -6 | Z'Lyte |
| RPS6KA1 (RSK1) | -1 | Z'Lyte |
| RPS6KA2 (RSK3) | 0 | Z'Lyte |
| RPS6KA3 (RSK2) | 10 | Z'Lyte |
| RPS6KA4 (MSK2) | -4 | Z'Lyte |
| RPS6KA5 (MSK1) | 2 | Z'Lyte |
| RPS6KA6 (RSK4) | 4 | Z'Lyte |
| RPS6KB1 (p70S6K) | 3 | Z'Lyte |
| RPS6KB2 (p70S6Kb) | -4 | Z'Lyte |
| SBK1 | 6 | Z'Lyte |
| SGK (SGK1) | 4 | Z'Lyte |
| SGK2 | 3 | Z'Lyte |
| SGKL (SGK3) | 0 | Z'Lyte |
| SNF1LK2 | -5 | Z'Lyte |
| SRC N1 | 5 | Z'Lyte |
| SRC | 14 | Z'Lyte |
| SRMS (Srm) | 2 | Z'Lyte |
| SRPK1 | 1 | Z'Lyte |
| SRPK2 | -4 | Z'Lyte |
| STK22B (TSSK2) | -2 | Z'Lyte |
| STK22D (TSSK1) | 6 | Z'Lyte |
| STK23 (MSSK1) | -2 | Z'Lyte |
| STK24 (MST3) | 4 | Z'Lyte |
| STK25 (YSK1) | -7 | Z'Lyte |
| STK3 (MST2) | -2 | Z'Lyte |
| STK4 (MST1) | -5 | Z'Lyte |
| SYK | 1 | Z'Lyte |
| TAOK2 (TAO1) | -1 | Z'Lyte |
| TBK1 | 4 | Z'Lyte |
| TEK (TIE2) Y897S | 2 | Z'Lyte |
| TEK (Tie2) | 5 | Z'Lyte |
| TNK1 | -3 | Z'Lyte |
| TXK | 4 | Z'Lyte |
| TYK2 | 0 | Z'Lyte |
| TYRO3 (RSE) | -5 | Z'Lyte |
| YES1 | -6 | Z'Lyte |
| ZAP70 | 4 | Z'Lyte |

**Figure S1.**
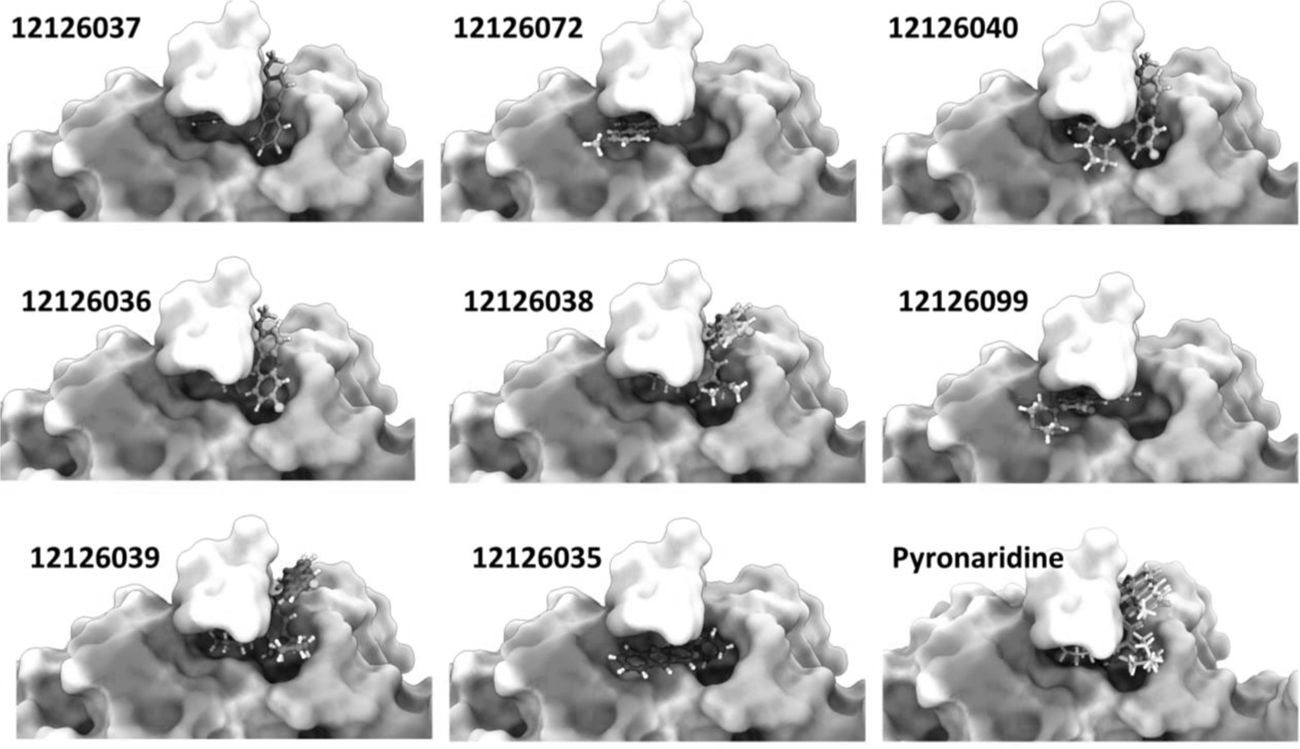
Docking poses from pyronaridine and all tested analogs. PL^pro^ surface is colored according to its electrostatic potential.

